# Design and Mathematical Analysis of Activating Amplifiers that Enable Modular Temporal Control in Synthetic Circuits

**DOI:** 10.1101/2023.03.15.532861

**Authors:** Calvin Lam

## Abstract

The ability to control mammalian cells such that they self-organize or enact therapeutic effects as desired has incredible implications. Not only would it further our understanding of native processes such as development and the immune response, but it would also have powerful applications in medical fields such as regenerative medicine and immunotherapy. This control is typically obtained by synthetic circuits that use synthetic receptors, but control remains incomplete. For example, the synthetic juxtacrine receptors (SJRs) are widely used as they are fully modular and enable spatial control, but they have limited gene expression amplification and temporal control. I therefore designed transcription factor based amplifiers that amplify gene expression and enable unidirectional temporal control by prolonging duration of target gene expression. Using an *in silico* framework for SJR signaling, I combined these amplifiers with SJRs and show that these SJR amplifier circuits can improve the quality of self-organization and direct different spatiotemporal patterning. I then show that these circuits can improve chimeric antigen receptor (CAR) T cell tumor killing against heterogenous and homogenous antigen expression tumors. These amplifiers are flexible tools that improve control over SJR based circuits and have both basic and therapeutic applications.

**Graphical Abstract:** 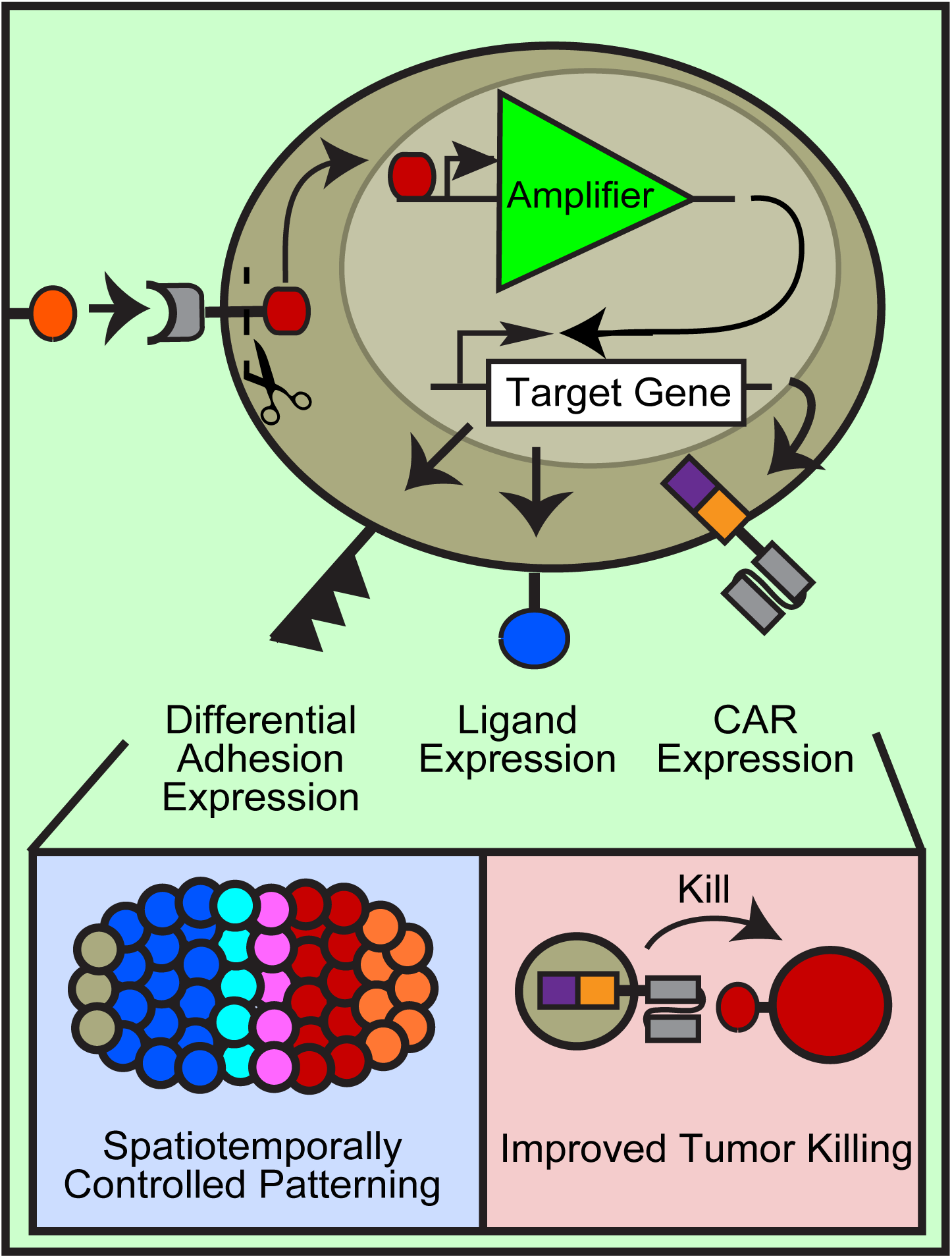

## INTRODUCTION

Although an emerging field, synthetic biology has already begun branching into two notable subfields: synthetic development and synthetic immunotherapy ^1–6^. In synthetic development, researchers are engineering mammalian cells to control self-organization; the goal is to further understanding of native developmental processes and regenerative therapeutics ^1, 4, 5, 7, 8^. In a referential study, naïve fibroblasts were programmed with synthetic circuits such that individual cells could sense neighbors and self-organize into diversely patterned spheroids ^9^. In synthetic immunotherapy, researchers focus on engineering immune cells to control therapeutic effects; the goal is to direct specific and novel immune responses ^2, 3, 10–14^. Recent achievements include programming T cells with synthetic circuits that enable multi-antigen discrimination, localized activation, and/or select cytokine secretion ^15–18^.

Both subfields commonly rely on synthetic receptors to create the behavior-controlling synthetic circuits, with synthetic juxtacrine receptors (SJRs) such as synNotch and SNIPRs being widely used due to their remarkable modularity, high spatial control, and potential for clinical translation ^9, 15–22^. The SJ class of receptors have an extracellular domain that can be programmed to sense a ligand of choice and the intracellular domain programmed to control a gene of choice (Fig.1A). When the SJR binds the juxtacrine target ligand on a neighboring surface, it releases an intracellular transcription factor to regulate the target gene ^16, 17, 19^ (Fig.1B). Because of this stoichiometric mechanism, one transcription factor released per activated SJR, there is minimal amplification of target gene expression ^19^.

**Figure 1.**
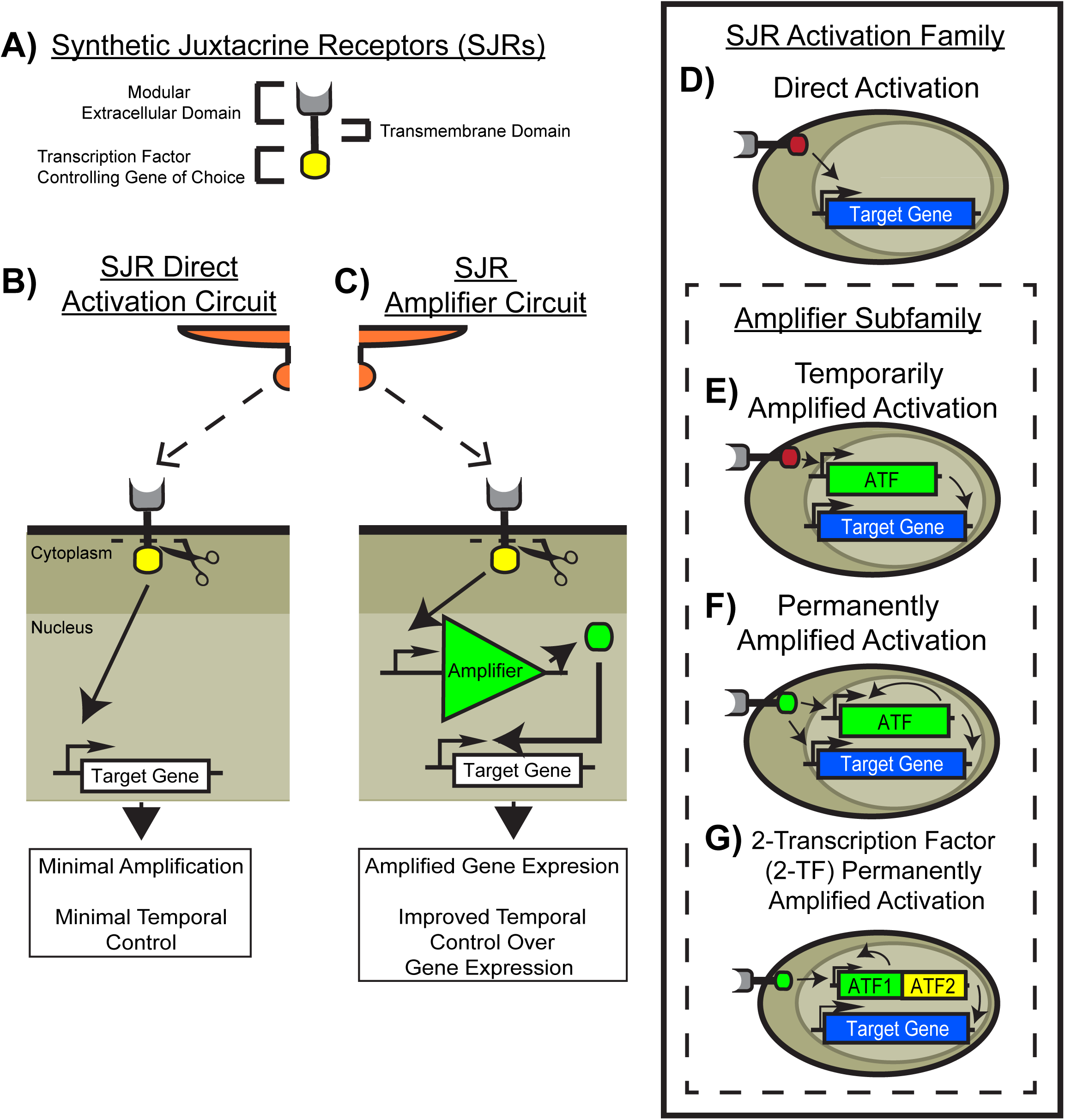
Designing Amplifiers to Combine with SJRs for the SJR Amplifier Circuits. A) Synthetic juxtacrine receptors (SJRs) are a class of synthetic receptors modular in both extracellular domain and intracellular domain. The extracellular domain consists of a choice ligand binding domain, and the intracellular domain consists of a transcription factor controlling a gene of choice ^9, 15–21, 35, 36^. B) In the classic SJR direct activation circuit, the SJR binds its juxtacrine cognate ligand to release the transcription factor controlling the target gene. This stoichiometric mechanism results in minimal expression amplification and temporal regulation ^16, 19^. C) This could be overcome by having the SJR drive the expression of a transcription factor based amplifier that amplifies target gene expression and prolongs duration of expression. D) The SJR Activation Family of circuits investigated in this study, consisting of the stereotypical direct activation circuit and the amplifier imbued Amplifier Subfamily. E) The temporarily amplified activation circuit is designed to temporarily amplify and prolong duration of target gene expression. SJR drives expression of an activating transcription factor (ATF) that then drives expression of the target gene. F) In the permanently amplified activation circuit, the SJR drives expression of an ATF that can then drive itself and the target gene, which should enable amplification and permanent gene expression. G) In the 2-transcription factor (2-TF) permanently amplified activation circuit, the SJR drives the expression of two different ATFs. ATF1 (green) then drives expression of itself and ATF2 (yellow). ATF2 then drives expression of the target gene.

Furthermore, as the SJR is dependent on juxtacrine ligand on neighbors to maintain gene expression, target gene expression is highly localized. This allows excellent spatial control over cell behavior, but minimal control on the temporal axis (Fig.1B) ^16, 19^. Thus, direct gene activation by SJRs neither enables amplification nor temporal regulation of target gene expression.

However, both expression amplification and temporal regulation are basic yet integral facets of numerous biological processes ^23–28^. For example, expression amplification of Nanog and Oct4 via a feedback loop is critical for embryonic stem cell maintenance ^27^. In the immune response to tumors, localized reservoirs of IL-2 serve to temporally enhance T cell activity ^28^. Somite development in vertebrates is an intricate dance involving both amplification and temporal regulation ^23–26^. As the goal of the subfields is to control such processes, it would be ideal if SJRs allowed for amplification and temporal regulation. Although the SJR’s current use form, the direct activation circuit, neither enables amplification nor temporal regulation, it may still be possible through an intermediate component ^16^.

Across diverse native biological circuits, transcription factors are used to regulate one another and the target gene in an amplifier manner to achieve temporal regulation ^27, 29–31^. In synthetic circuits, transcription factors have been used to generate amplifier loops, allowing cells to retain memory of a stimulus ^32^. As SJRs mechanistically function via transcription factors, I reasoned that a transcription factor based amplifier could amplify target gene expression and concurrently enable temporal regulation in SJR based circuits.

I therefore designed several transcription factor based amplifiers that enable expression amplification and temporal regulation and combined them with SJRs. These SJR amplifier circuits use a SJR to drive expression of an amplifier gene circuit that subsequently regulates target gene expression (Fig.1C). In this amplifier subfamily, which amplifies gene expression and increases duration of gene expression, I designed three circuits. All three amplify gene expression but one temporarily prolongs gene expression (Temporarily Amplified Activation, Fig.1E) and two enable permanent gene expression (Permanently Amplified Activation Fig.1F and 2-TF Permanently Amplified Activation Fig.1G). The SJR inhibition family and its amplifier variants, designed to amplify inhibition of gene expression and decrease duration of gene expression, is described in the upcoming accompanying paper.

I chose to study these circuits with an *in silico* approach as it would enable rapid yet extensive investigation into circuit behavior across a variety of conditions. I used the framework by Lam et al. as it has been shown to faithfully model SJR circuits via the GJSM model and successfully predict self-organization in SJRs driving differential adhesion ^33^. Using the framework, I converted the circuits into mathematical equations, then implemented the equations into *in silico* cells created in the cellular Potts model ^26, 33, 34^. This is the *in silico* analog of transducing live cells with analogous circuit genes (SFig.1A).

Here I show that the amplifiers are capable of amplifying target gene expression and increasing duration of gene expression. When combined with SJRs, these SJR amplifier circuits enable improving self-organization and directing novel spatiotemporal patterning in synthetic development. In synthetic immunotherapy, these circuits improve tumor cell killing. Though I focus here on two subfields, these amplifiers serve as versatile tools to temporally control and amplify gene expression for a broad range of synthetic biology applications.

## RESULTS

### Design of the SJR amplifier circuits

Of the circuits in the SJR Activation Family, the direct activation circuit is the most well-known. It uses a synthetic juxtacrine receptor (SJR) that, upon binding its cognate ligand, releases a transcription factor that directly drives target gene expression (Fig.1D) ^9, 15–21, 35, 36^. This design offers high spatial control, but with strong reliance on SJR signaling to maintain target gene expression, it offers littl temporal control. Furthermore, due to SJR stoichiometry (one transcription factor (TF) per SJR), it offers little target gene expression amplification ^16, 19^.

The Amplifier Subfamily is designed to add gene expression amplification along with temporal control to SJR based circuits (Fig.1E-G). In the temporarily amplified activation circuit, the SJR drives expression of an activating transcription factor (ATF) that subsequently drives expression of the target gene (Fig.1E). Gene expression amplification should be observed with higher gene product levels.

Furthermore, continued increasing target gene expression should be observed even when SJR signaling is lost, provided that the ATF level remains sufficiently elevated to continue driving target gene expression. Thus, this temporarily amplified activation circuit should amplify and temporarily prolong target gene expression.

Amplification can also result in permanent gene expression, and I designed two variants to achieve this. In the permanently amplified activation circuit, the transcription factor released by the SJR is identical to the ATF. Furthermore, both the ATF gene and target gene are activated by this ATF (controlled by same type of promoter) (Fig.1F). Thus, with initial sufficient SJR signaling, ATF is expressed and drives ATF expression. As the target gene’s promoter is responsive to ATF, target gene expression should be amplified and permanently driven.

In the 2-transcription factor (2-TF) permanently amplified activation circuit, the SJR drives the expression of two different ATFs, ATF1 and ATF2. These two ATFs are under the same promoter and are on the same transgene cassette. ATF1 is identical to the SJR’s transcription factor thus as in the permanently amplified activation circuit, ATF1 drives ATF1 expression. As ATF2 is in the same cassette as ATF1, it becomes driven, amplified, and permanently expressed. The target gene is controlled by a different type of promoter responsive to only ATF2 and thus, as ATF2 becomes amplified and permanently expressed, so does the target gene (Fig.1G).

### Amplifiers amplify and enable different levels of temporal control over target gene expression

With the SJR amplifier circuits designed, I converted the circuits to equations using the GJSM model ^33^ and implemented them into *in silico* cells defined via the Cellular Potts framework ^26, 34^ (See Methods section for more details). I then tested their ability to amplify target gene expression and enable temporal control with a simple cell-cell signaling setup modified from ^37^. Gray cells were programmed with either the direct activation circuit (Fig.1D) or one of the amplifier circuits (Fig.1E-G), with the SJR set to be responsive to orange ligand on orange cells and the target gene set to be a red reporter (Fig.2A).

**Figure 2.**
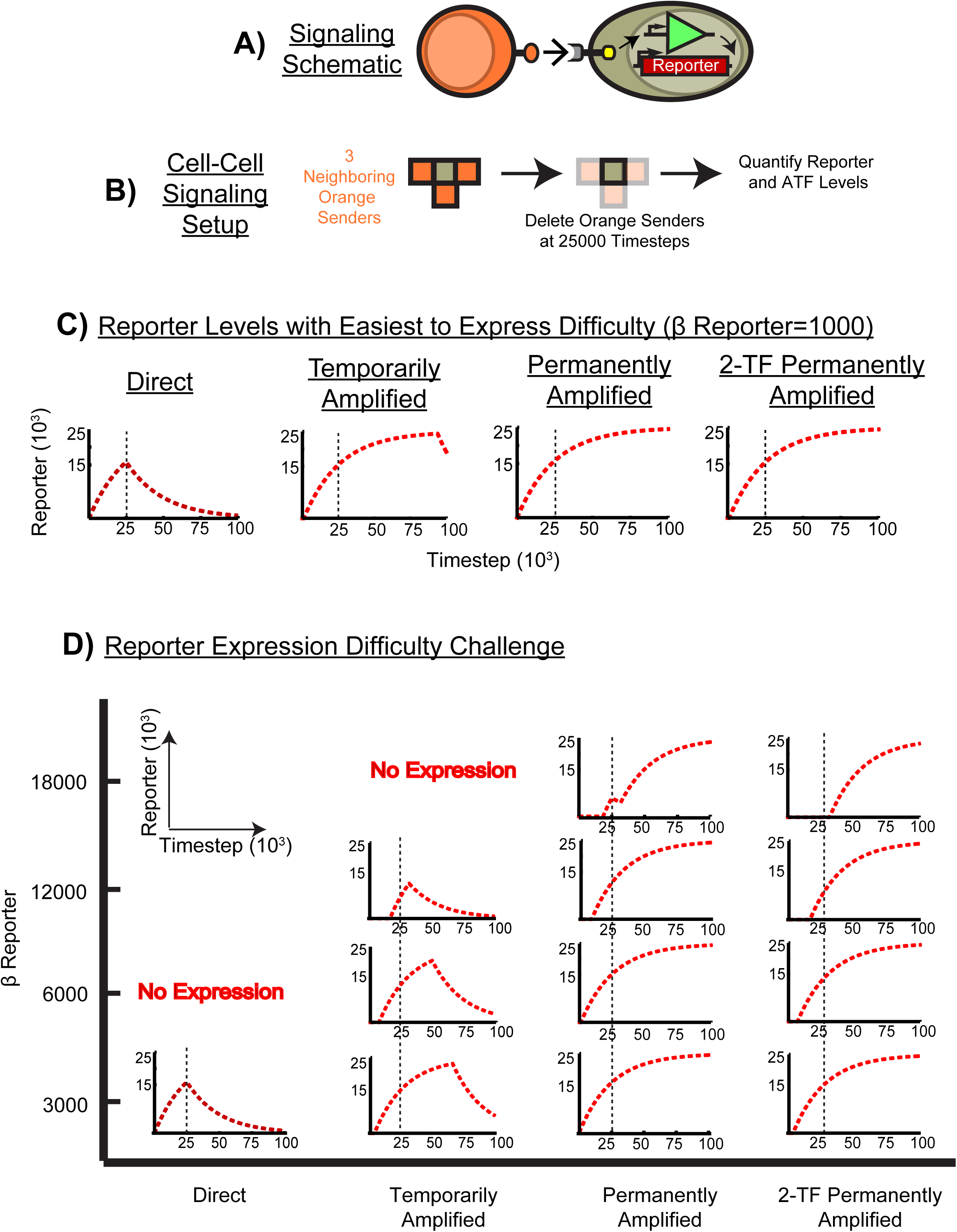
Amplifiers Amplify and Enable Temporal Regulation Over Target Gene Expression. A) Signaling schematic used to test the amplifiers for their ability to amplify gene expression and enable temporal control of a red reporter. Gray cells receive signal from orange ligand on orange cells via an anti-(orange ligand) SJR that either drives red reporter expression directly (Direct Activation) or an amplifier (Temporarily Amplified, Permanently Amplified, or 2-TF Permanently Amplified Activation) that then drives red reporter expression. B) One cubic gray cell is seeded with 3 cubic orange cells and at 25,000 timesteps, orange cells are deleted to determine effect on red reporter levels. Both reporter and ATF levels were tracked. Amplifiers should amplify reporter levels such that there should be continued increasing reporter levels immediately after SJR signal loss and higher reporter levels. C) Reporter traces from the different circuits. At the easiest reporter expression difficulty (β Reporter=1000), directly activated cells had reporter expression decrease immediately after loss of orange cells while the amplifier imbued cells continued to increase reporter levels and maintain higher reporter levels. D) Reporter expression difficulty challenge. Reporter levels from challenging the cells with higher reporter expression difficulty, β Reporter, from 3000 to 18000. At the same expression difficulty, amplifier imbued cells had increasing reporter levels even after orange neighbors were lost and higher reporter levels, indicating amplification of target gene expression. These results also confirm the temporal design of circuits. The temporarily amplified activation circuit only temporarily prolongs reporter expression (decreases sometime later) while the two permanently amplified circuits continue to prolong reporter expression. One trace shown per circuit/condition and simulations ran for 100,000 timesteps.

A single cubic gray cell was then seeded with 3 orange cubic cells such that one face of an orange cell contacted one gray cell face (Fig.2B). Cells were frozen (static in volume, surface area, and morphology) to maintain uniform and constant SJR signaling. This cell-cell signaling setup enables controlling the exact orange ligand-SJR signal the gray cell receives and should therefore enable determining the amplificatory effects of the amplifiers. At 25,000 timesteps, orange neighbors were deleted to test how SJR signaling loss would affect reporter expression (Fig.2B). Both reporter and ATF levels (if applicable), were tracked throughout the course of the experiment.

In the GJSM model, different properties of gene expression are modelled by different parameters. With a circuit that activates gene expression, the parameter β models difficulty of gene expression, with higher values modeling higher expression difficulty. Lower β values allow easier gene expression but at the risk of leaky gene expression ^33^. See Brief overview of Generalized Juxtacrine Signaling Model (GJSM) in the Methods Section or the original study ^33^ for more details.

With the reporter expression difficulty (β Reporter) set to 1000, the easiest tested expression difficulty without leakiness (SFig.2A), all circuits were able to yield reporter expression with 3 orange neighbors. Consistent with previous *in vitro* studies, the direct activation circuit had immediate reporter level decrease upon removal of the neighbors (Fig.2C) ^16, 19^. In contrast, all of the amplifier circuits were able to maintain increasing reporter levels for some time after loss of SJR signaling (Fig.2C). Moreover, these amplifier circuits were able to maintain higher reporter levels as well.

Challenging these circuits with higher reporter expression difficulties yielded similar results Amplifier circuits did not have immediate reporter decrease upon loss of SJR signaling and at the same reporter expression difficulty level, provided that the reporter was expressed, consistently had higher reporter levels compared to the direct activation circuit (Fig.2D). Similar results were obtained with 1 and 6 orange neighbors (data not shown). Therefore, with the same identical SJR signal that the direct activation circuit had, the amplifier circuits were able to amplify reporter expression to maintain higher reporter levels and prolong reporter expression.

Furthermore, these results confirm that the amplifier circuits operate temporally as designed. The temporarily amplified activation circuit only temporarily prolonged reporter expression, as reporter levels eventually decreased at all the reporter expression difficulties tested (Fig.2C-D). This is due to the ATF’s dependency on SJR signaling to remain elevated and sustain reporter expression (SFig.2B). In contrast, both the permanently amplified activation circuit and 2-TF permanently amplified activation circuit demonstrated no notable decrease in reporter levels, confirming their capability to permanently drive target gene expression (Fig.2C-D). This is due to the ATF being able to maintain elevated ATF levels via positive self-feedback (SFig.2B).

### Amplifiers improve 3-layered structure formation and enable temporal control in self-organization

With evidence that the amplifiers are able to amplify target gene expression and enable temporal control, I next tested these circuits in synthetic development experiments as the direct activation circuit has been used extensively to drive self-organization ^9^. In the reference study, naïve L929 mouse fibroblast cells were engineered with a variety of different direct activation circuits driving the expression of different cadherins, ligands, and reporters. As cells signaled to one another and sorted, they formed a variety of patterned spheroids. Of these spheroids, the 3-layered structure is the most referential; the robustness of its self-organization has been investigated both *in vitro* and *in silico* ^9, 33, 38, 39^. I therefore tested if the amplifier circuits could improve the self-organization of the 3-layered structure.

To achieve the 3-layered structure, I used the developmental trajectory in the original study ^9^. From a mixture of orange and gray cells, gray cells receive signal from orange cells and become high E-cadherin positive (blue/cyan colored cells) (Fig.3A). These high E-cadherin cells form a core and signal back to orange cells, turning orange cells low E-cadherin positive (red colored). End result is a 3-layered structure with a high E-cadherin blue/cyan core surrounded by a low E-cadherin red ring which is surrounded by unactivated orange cells (Fig.3A). The color key to the cell’s state is given in Fig.3B. Ecad is E-cadherin and ATF is activating transcription factor.

**Figure 3.**
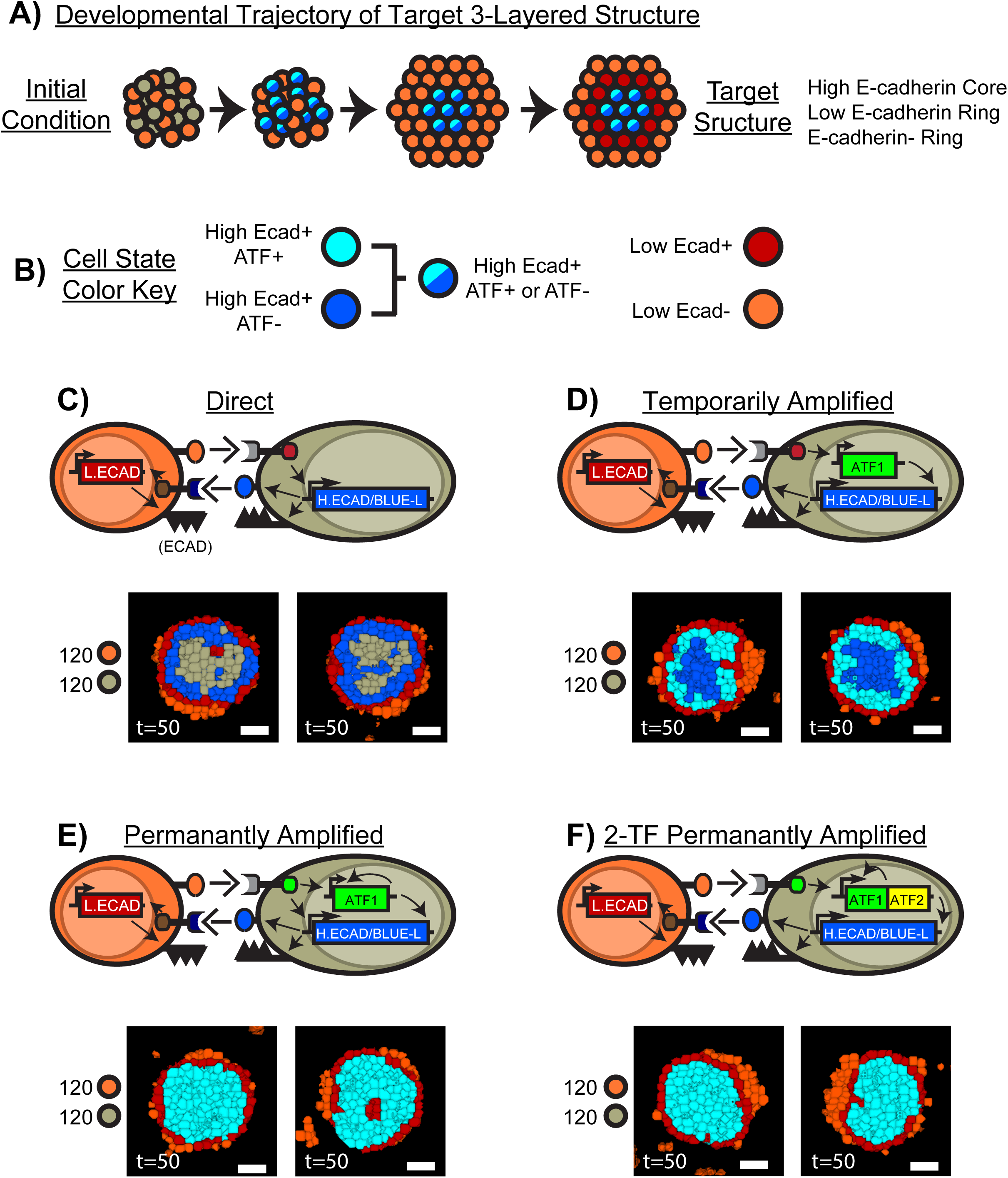
Amplifiers Improve 3-Layered Structure Formation. A) Amplifiers were tested if their amplification and extension of gene expression duration could improve 3-layered structure formation. In the original 3-layered structure developmental trajectory ^9^, a mixture of gray and orange cells is seeded. Orange cells signal to gray cells and turn them high E-cadherin^+^ (blue or cyan). These blue/cyan cells express a ligand that signals back to orange cells to turn them low E-cadherin^+^ (red). This should result in a 3-layered structure with a blue/cyan core followed by a red ring followed by an orange ring. B) Cell state color key is given, describing what color corresponds to the expression state of the cell. Ecad is E-cadherin and ATF is activating transcription factor. C) Representative cross sections of two 3-layered structure formed from directly activated orange cells and gray cells. Circuits are given. Anti-(orange ligand) SJR on gray cells directly drives high E-cadherin and blue ligand expression but consistently results in poor structure formation with the core marred by high E-cadherin^-^ gray cells. D) Cross sections of two 3-layered structure formed from directly activated orange cells and temporarily amplified activated gray cells. The amplifier driving high E-cadherin and blue ligand expression robustly improves 3-layered structure formation, yielding a mixed blue and cyan core of high E-cadherin^+^ cells. E) Permanently amplified activated gray cells also strongly improve 3-layered structure formation but yields a completely cyan core instead. F) 2-TF Permanently amplified activated gray cells improve 3-layered structure formation with a cyan core as well. Mixtures are of 123.8±2.67 orange cells and 127.2±2.67 gray cells. N=5 for each with two representative cross sections shown. Simulations run for 50,000 timesteps. Additional structures and results are given in SFig.3.

In the original direct activation circuit, cell-cell signaling is achieved via SJRs ^9^. Gray cells express an SJR that, in response to orange ligand on orange cells, induces high E-cadherin (gene is *H.ECAD*) along with blue ligand (gene is *BLUE-L*) expression (Fig.3C). With blue ligand on their surface, gray cells turn blue and signal to orange cells as orange cells constitutively have anti-(blue ligand) SJR on their surface. This induces low E-cadherin (gene is *L.ECAD*), turning orange cells red (Fig.3C). I programmed this direct activation circuit into *in silico* L929 (ISL929), an *in silico* cell line modelling the same murine L929s used in the original study and validated for modeling such structures ^9, 33^. However, despite using a favorable ratio (∼120 gray to ∼120 orange) and making high E-cadherin and blue ligand easy to express (β Adhesion/Ligand=1000), this direct activation circuit consistently (5/5) resulted in poor 3-layered structure formation (Fig.3C). Although the red ring and orange ring were obtained, the desired high E-cadherin core was consistently marred by E-cadherin^-^ gray cells.

Quantifying core quality with a homogeneity index ^33^ that measures high E-cadherin^+^ cell connectedness (i.e. high E-cadherin^+^ cells contact to other high E-cadherin^+^ cells, see Homogeneity Index in the Methods Section or the original study ^33^ for more details), confirmed this observation; core quality saturated around ∼0.7 while similar reference structures typically exceed 0.8 (Core Quality, SFig.3A) ^33^. This was not due to gray cells being unable to become high E-cadherin^+^ blue cells. The majority of gray cells became blue as shown by the developmental trajectory and the relative activation plot, but they revert to gray cells as time passed (Developmental Trajectory and Relative Activation, SFig.3A). This is consistent with SJR direct activation to be minimally amplificatory and highly contact dependent ^16, 19^.

Challenging these cells with different ratios that used less orange cells resulted in even poorer structure formation, again off target from the targeted 3-layer structure (Different Ratios, SFig.3A). Similarly poor 3-layered structures formed with the tested ratios at a higher high E-cadherin and blue ligand expression difficulty (β Adhesion/Ligand=3000, data not shown).

Using the temporarily amplified activation circuit instead strongly improved target structure formation, consistently yielding 3-layered structure formation in all 5/5 replicates (Fig.3D). An example developmental trajectory is given in SFig.3B. Although the core was a mix of blue (high E-cadherin^+^ ATF^-^) and cyan (high E-cadherin^+^ ATF^+^) cells, it consisted of all high E-cadherin^+^ cells as desired, indicating that high E-cadherin expression was prolonged (Fig.3D). Moreover, more gray cells became blue/cyan (high E-cadherin^+^) compared to the direct activation circuit (blue cell curve+cyan cell curve ∼100% vs blue cell curve ∼85% at peak activation at around 20,000 timesteps), indicating high E-cadherin expression was amplified (Relative Activation, SFig.3B). Core quality exceeded that in the direct activation circuit and is similar to the values of other quality 3-layered structures (Core Quality, SFig.3B) ^33^. Furthermore, at ratios with less orange cells, gray cells with the temporarily amplified activation circuit still managed to become high E-cadherin^+^ (blue or cyan), allowing target core formation as desired. However, at these ratios the red and orange ring became diminished as is observed *in vitro* (Different Ratios, SFig.3B) ^9^. Similar structures were obtained with the tested ratios with β Adhesion/Ligand=3000 (data not shown).

The cell-cell signaling experiment (Fig.2) showed that the temporarily amplified activation circuit only temporarily prolongs target gene expression (Fig.2C-D). It is therefore expected that core obtained with the temporarily amplified activation circuit will eventually degenerate into a blue/cyan core marred with gray cells like those obtained with the direct activation circuit. Indeed, running the experiment to 75,000 timesteps results in blue cells beginning to deactivate to E-cadherin^-^ gray cells (Deactivation Test, SFig.3B). This can be circumvented, if desired, by amplifying and indefinitely prolonging high E-cadherin and blue ligand expression with one of the permanent amplification circuits (Fig.3E-F).

Both circuits yielded similar results: robust 3-layered structure formation (5/5 replicates for both) with cores of cyan (high E-cadherin^+^ ATF^+^) cells (Fig.3E-F). Core quality reflected this, exceeding 0.8 (Core Quality, SFig.3C-D). Compared to the direct activation circuit, more cells became high E-cadherin^+^ and none reverted from being high E-cadherin^+^ (Relative Activation, SFig.3C-D). These results indicate that high E-cadherin expression was both amplified and prolonged. Given that the temporarily amplified activation circuit was able to yield high E-cadherin^+^ (blue or cyan) cores when challenged with lower orange cell ratios, it is unsurprising that these two circuits were also able to yield high E-cadherin^+^ target cores (Different Ratios, SFig.3C-D). Developmental trajectory is given in SFig.3C and D. Similar structures were obtained with the tested ratios with β Adhesion/Ligand=3000 (data not shown).

These data indicate that the amplifiers are able to improve self-organization; amplifying and prolonging high E-cadherin/blue ligand expression robustly improved target 3-layered structure formation. Moreover, these amplifiers enable temporal control over self-organization: the temporarily amplified activation circuit allows generating a temporary 3-layered structure while the permanently amplified circuits allow a permanent 3-layered structure.

### Spatial control from SJRs and temporal control from amplifiers enable spatiotemporal patterning

Combining the amplifiers with the SJRs yielded SJR amplifier circuits that allowed robust formation of the target 3-layered structure, but there were noticeable differences in the spatial patterning between the temporarily amplified activation circuit and the permanent amplification circuits. Due to differences in temporal control, the temporarily amplified activation circuit formed a core consisting of blue cells with a cyan ring (Fig.3D) while the permanent amplification circuits formed cores with all cyan cells (Fig.3E-F). This suggests that the SJR amplifier circuits can exert spatiotemporal control over multicellular patterning.

To confirm this, I tested the amplifiers on another structure, non-concentric and asymmetrical multipole, that is also from the reference study ^9^. The signaling steps are like those used in the 3-layered structure, but the signaling instead drives the expression of two different, but highly homotypic adhesion proteins N-cadherin and P-cadherin. Gray cells receive signal from orange cells and become N-cadherin (Ncad) positive with signaling capability to orange cells (Fig.4A). In response, orange cells express P-cadherin (Pcad), turn red, and adhere to one another (Fig.4A). A mixture of these gray and orange cells results in a multipole structure with cores/poles of either N-cadherin^+^ (Ncad^+^) or P-cadherin^+^ (Pcad^+^) cells (Fig.4B).

**Figure 4.**
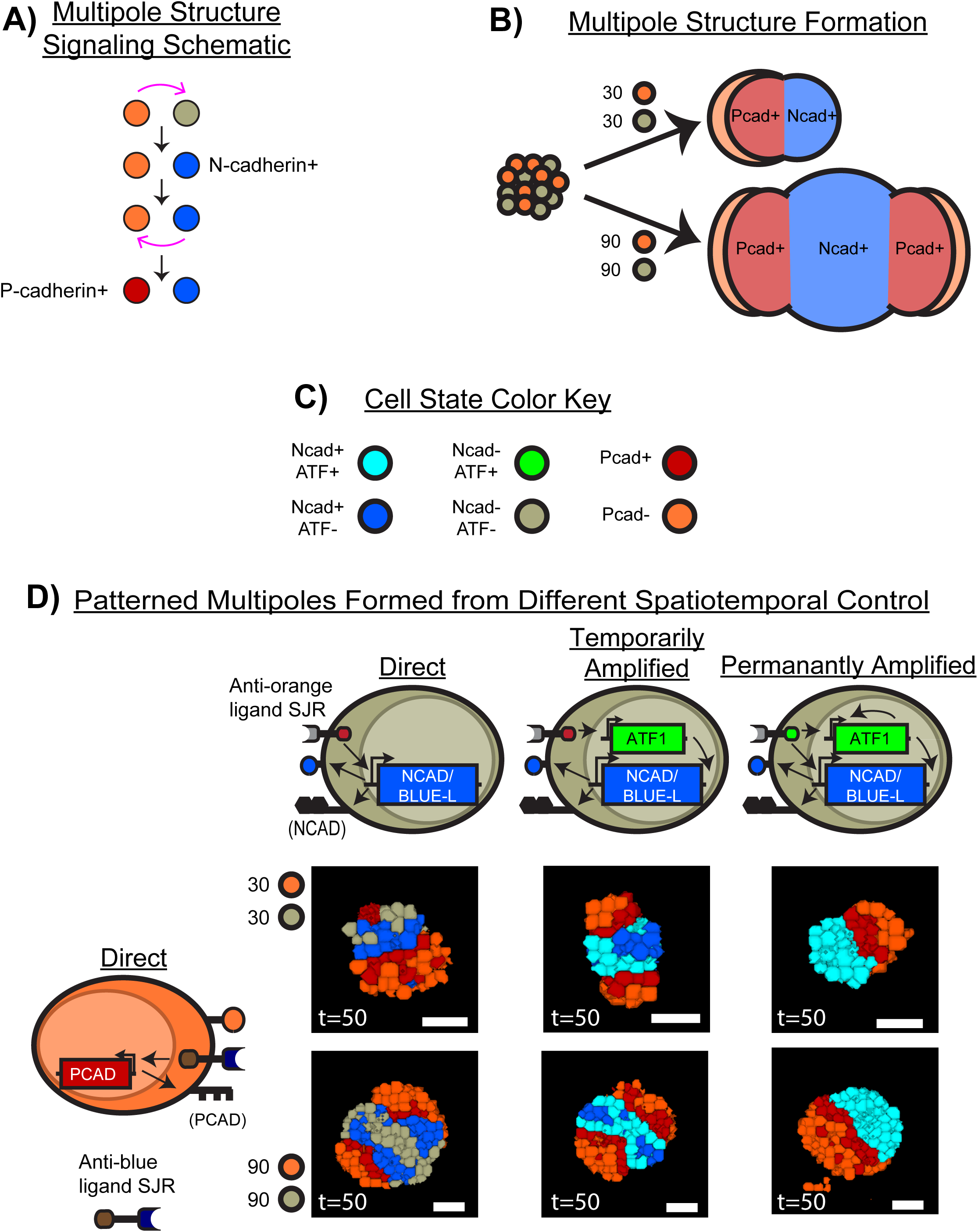
Combining Amplifiers with SJRs Creates SJR Amplifier Circuits That Enable Spatiotemporal Patterning. A) SJR amplifier circuits were tested for the ability to direct spatiotemporal patterning in a multipole structure. In the original multipole structure schematic, orange cells signal to gray cells turning them blue N-cadherin^+^. Blue N-cadherin^+^ cells express a ligand enabling them to signal back to orange cells, turning them into red P-cadherin^+^ cells. B) N-cadherin and P-cadherin are homotypically adhesive ^9, 33^, thus mixing orange and gray cells results in multipole structures. C) Cell state color key is given, describing what color corresponds to the expression state of the cell. Ncad is N-cadherin, Pcad is P-cadherin, ATF is activating transcription factor. D) Different circuit combinations and a cross section of the resulting multipole structure. Orange cells always have the direct activation circuit, with P-cadherin (gene is *PCAD*) expression driven by an anti-(blue ligand) SJR. Gray cells have different activation circuits; the anti-(orange ligand) SJR either drives N-cadherin and blue ligand (gene is *NCAD/BLUE-L*) expression directly (Direct Activation) or an amplifier (Temporarily Amplified or Permanently Amplified) that then drives N-cadherin and blue ligand expression. Combining these orange and gray cells demonstrates that different amplifiers enable different temporal control that dictates spatiotemporal patterning. Directly activated gray cells characteristically form a blue stripe with a gray core/pole. Temporarily amplified activated gray cells amplify and prolong N-cadherin expression to form a cyan stripe with a blue core/pole. Permanently amplified activated gray cells permanently amplify and prolong N-cadherin expression to form cyan cores/poles. Mixtures are of 27.4±1.04 orange cells with 29.6±1.04 gray cells or 87.4±1.47 orange cells with 91.6±1.47 gray cells. N=10 for each circuit combination with one representative cross section shown. Simulations run for 50,000 timesteps. Additional cross sections are given in SFig.4.

I therefore programmed gray cells with either the direct or an amplifier circuit. The 2-TF permanently amplified activation circuit was not tested due to its patterning similarity to the permanently amplified activation circuit (Fig.3E-F, SFig.3C-D). The SJR on gray cells responds to orange ligand on orange cells and drives either N-cadherin/blue ligand (gene is *NCAD/BLUE-L*) directly (direct activation circuit) or an amplifier that subsequently drives N-cadherin/blue ligand (amplifier circuits). This turns gray cells blue or cyan, representing the N-cadherin^+^ blue ligand^+^ state. Blue/cyan cells can now signal via blue ligand to orange cells, as they constitutively have anti-(blue ligand) SJR on their surface. This directly drives P-cadherin (gene is *PCAD*) expression in orange cells. The color key to the cells’ states is given in Fig.4C. Ncad is N-cadherin, Pcad is P-cadherin and ATF is activating transcription factor. These described circuit combinations are given in Fig.4D.

Combining the orange cells with differently activated gray cells resulted in a variety of spatiotemporal patterned structures. Directly activated gray cells consistently formed a spatially sensitive pattern; blue cells were localized where they could contact cognate signaling orange/red cells while cells that lost contact deactivated to gray, forming the rest of the core/pole (Fig.4D). A mixture of ∼30 gray and ∼30 orange cells yielded a blue stripe with a gray pole while ∼90 gray and ∼90 orange cells yielded a gray core with blue stripes proximal to orange/red cells (Fig.4D). Additional representative structures are given in SFig.4.

Adding temporal control with temporarily amplified activated gray cells generated a spatiotemporal pattern; cyan cells were localized next to orange/red cells but amplifying and prolonging high N-cadherin/blue ligand expression allowed cells to remain blue (N-cadherin^+^ blue ligand^+^) even when they lost contact with orange/red cells (Fig.4D). ∼30 gray and ∼30 orange cells generated cyan stripes with a blue pole while ∼90 gray and ∼90 orange cells formed a blue core with cyan stripes next to orange/red cells (Fig.4D). Additional representative structures are given in SFig.4.

Adding temporal control with the permanently amplified activation circuit yielded a different spatiotemporal pattern; nearly all gray cells became cyan (N-cadherin^+^ blue ligand^+^) and cyan cells localized independent of contact with orange/red cells (Fig.4D). These results are consistent with the permanently amplified activation circuit’s design; it amplifies and drives permanent expression of the target gene, *NCAD/BLUE-L*, thus loss of contact with orange/red cells should not affect target gene expression. ∼30 gray and ∼30 orange cells generated cyan poles while ∼90 gray and ∼90 orange cells reliably formed cyan cores (Fig.4D). Additional representative structures are given in SFig.4.

These results confirm the observations with the 3-layered structure. Different amplifiers confer different levels of temporal control and when combined with the spatial control conferred by SJRs, yield spatiotemporal circuits that can direct multicellular patterning.

### Spatiotemporal control from SJR amplifier circuits can be intercellularly compatible

In the multipole structure formation, orange cells are dependent on blue/cyan cell signaling (via blue ligand) to become P-cadherin^+^ (Fig.4A). Intriguingly, however, orange cells consistently formed orange poles with a red stripe regardless of the activation circuit used in gray cells (Fig.4D, SFig.4). This suggests that the amplifiers can have a minimal effect on the activation circuits in other cells, even if those activation circuits are dependent on the amplified gene for signaling. This is supported by the results with the 3-layered structures; the red and orange ring pattern was similar across all 3-layered structures despite gray cells having different activation circuits (Fig.3C-F). I therefore hypothesized that the SJR amplifier circuits can pattern independently of the activation circuits in cognate signaling cells.

To test this, I used the same setup for multipole structure formation (Fig.4), but also expanded orange cells to have either the temporarily amplified activation circuit or the permanently amplified activation circuit. This allowed me to test the remaining combinations of circuits not tested in Fig.4: gray (direct, temporarily amplified, permanently amplified) to orange (temporarily amplified, permanentl amplified) (Fig.5A). The color key to the cells’ states is given in Fig.5B. Pcad is P-cadherin, Ncad is N-cadherin and ATF is activating transcription factor.

**Figure 5.**
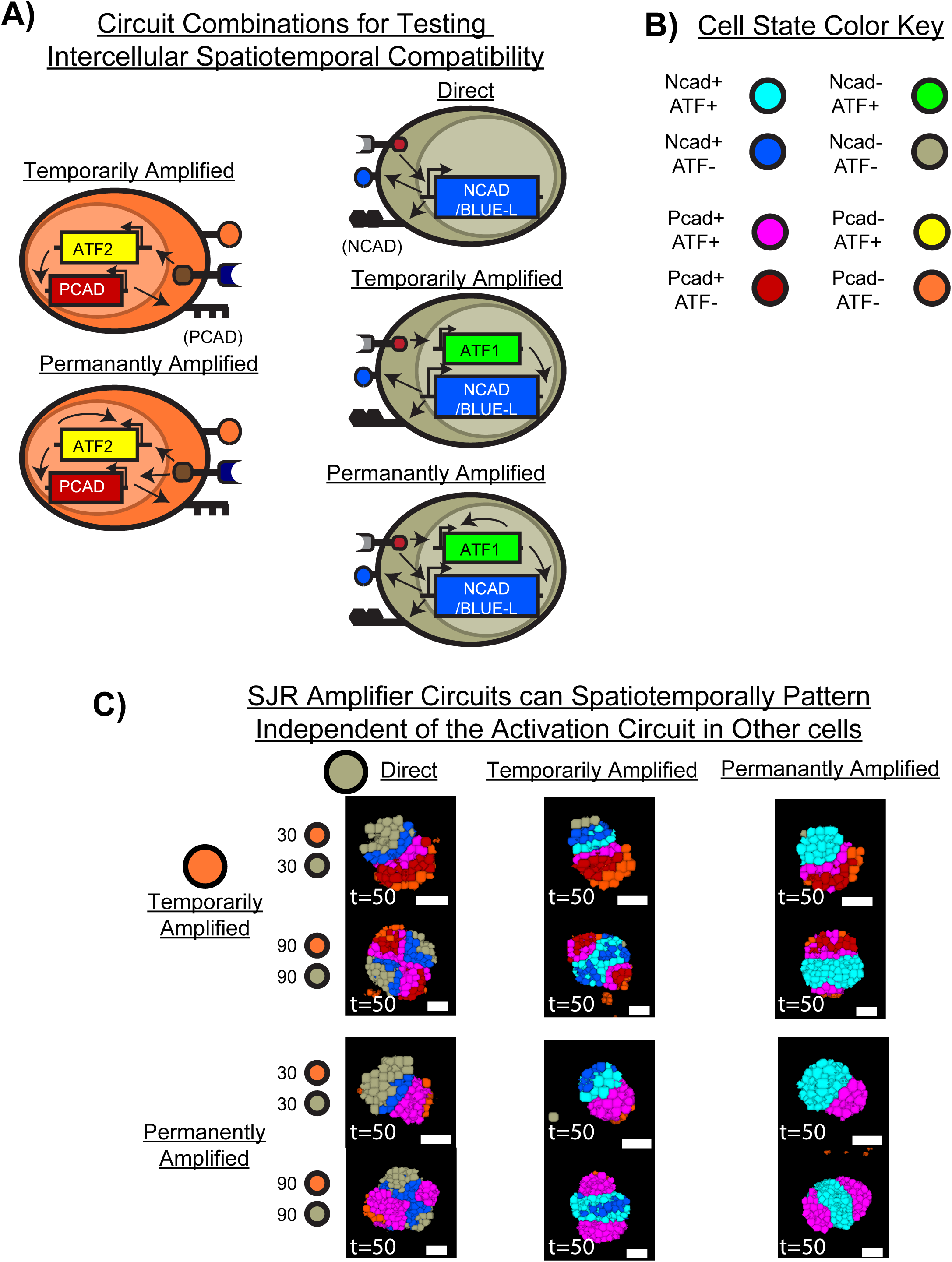
SJR Amplifier Circuits Enable Intercellularly Compatible Spatiotemporal Control. A) Orange cells were programmed with either the temporarily amplified activation or permanently amplified activation circuit, then mixed with differently activated gray cells to determine if orange cells could spatiotemporally pattern independent of the circuit in the gray cells. B) Cell state color key is given, describing what color corresponds to the expression state of the cell. Ncad is N-cadherin, Pcad is P-cadherin and ATF is activating transcription factor. C) Different circuit combinations and a cross section of the resulting multipole structure. Across two different cell amounts (27.4±1.04 orange cells with 29.6±1.04 gray cells or 87.4±1.47 orange cells with 91.6±1.47 gray cells), orange cells yielded patterning independent of the activation circuit in gray cells. Temporarily amplified activated orange cells consistently formed a pink, red, and orange striped core/pole. Permanently amplified activated orange cells consistently formed a pink core/pole. N=10 for each circuit combination with one representative cross section shown for each initial cell amount. Simulations run for 50,000 timesteps. Additional cross sections are given in SFig.5.

Orange cells are dependent on blue ligand from blue/cyan cells to become P-cadherin^+^ and red (Fig.5A), but their spatiotemporal patterning was not dependent on the amplification of blue ligand expression (Fig.5C). Temporarily amplified activated orange cells formed a pink, then red, and then orange stripe (Fig.5C). Permanently amplified activated orange cells formed a pink pole with several orange cells on the periphery (Fig.5C). These patterns were consistently obtained independent of the activation circuit in gray cells. Additional representative structures are given in SFig.5. This was further confirmed quantitatively; quality of the individual poles (blue/cyan poles and red/pink poles) as measured by the homogeneity index and activation of cells over time were independent of the circuit in the cognate signaling cell type (SFig.6). For pole quality, cyan curve is cyan/blue pole and pink curve is red/pink pole. For the relative activation plots, curve color corresponds to cell state color in Fig.5B.

**Figure 6.**
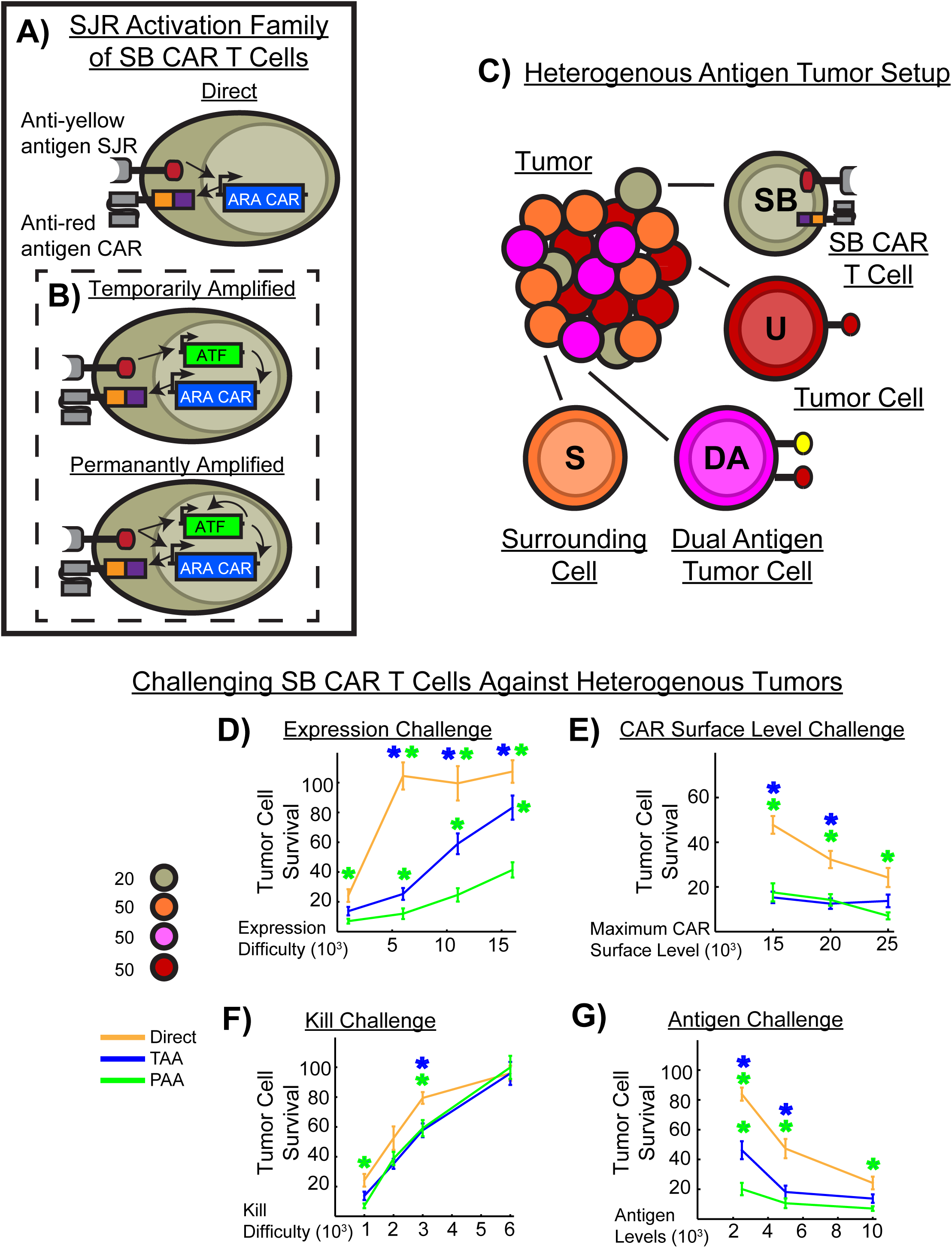
SJR Amplifier Circuits Improve CAR T Cell Killing Against Heterogenous Tumors Across a Variety of Conditions. A) Activation family of SB CAR T cells created by rewiring the cells used in the 3-layered and multipole structures. The SJR is rewired to sense anti-(yellow antigen) and the cadherin/blue ligand target gene is replaced with anti-(red antigen) CAR (*ARA CAR*). In the direct activation circuit, the SJR directly drives expression of ARA CAR. *In vitro*, these cells have highly localized anti-tumor effects and require both antigens to activate and kill tumor cells ^21, 36^. B) In the amplifier imbued SB CAR T cells, the SJR drives an amplifier that that then drives the ARA CAR. C) Creation of the heterogenous antigen tumor setup. These cells were tested against a heterogenous tumor *in vitro* ^21^ and to mimic this, I created the cells necessary to make an *in silico* model of this tumor. Orange surrounding S cells have no surface antigen and represent surrounding healthy cells. Pink dual antigen (DA) cells have yellow tumor antigen one and red tumor antigen two. Red tumor (U) cells have red tumor antigen two. Mixing these cells with the gray SB CAR T cells enables creating a variety of heterogenous tumors. Heterogeneous tumor here is 19.8±0.85 gray SB CAR T cells with 55.7±2.13 orange S cells with 52.5±2.33 red U cells and 51±1.82 pink DA cells. Circuit and curve color key is given to the left of the bottom plots. D) CAR Expression Challenge. SB CAR T cells were challenged with increasing difficulty of CAR expression (β CAR) and relative tumor cell survival quantified. Overall, using an amplifier to amplify and prolong CAR expression decreased tumor cell survival more than the direct activation circuit did. E) CAR Surface Level Challenge. SB CAR T cells were challenged with different maximum CAR surface levels (κ) and tumor cell survival quantified. At lower maximum CAR surface levels, SB CAR T cells with amplifiers decreased tumor cell survival more than the directly activated cells did. F) Kill Challenge. SB CAR T cells were challenged to kill increasingly CAR signaling resistant tumor cells. Amplifier imbued SB CAR T cells moderately decreased tumor cell survival more than the direct activated cells did, indicating that amplifying and prolonging CAR expression could aid in removing CAR signaling resistant tumors cells. G) Antigen Challenge. SB CAR T cells were challenged to kill lower antigen tumor cells. Amplifier imbued SB CAR T cells could amplify the smaller signal and kill more tumor cells. N=10 for each circuit for each parameter tested. * denotes significant difference and is colored the curve it differs from. Simulations run for 50,000 timesteps with shown data from this endpoint.

These data indicate that the SJR amplifier circuits can be used to pattern independent of the activation circuit in other cells, even if the amplifier circuits are dependent on signaling from those activation circuits. Therefore, different spatiotemporal control from different amplifier circuits can be modularly combined on the intercellular level to multiplex and expand the variety of spatiotemporal structures.

### SJR amplifier circuits improve CAR T cell tumor killing against heterogenous tumors

With evidence that the amplifiers amplify and enable temporal control over gene expression, and when combined with SJRs, yield circuits that improve self-organization and enable spatiotemporal patterning in synthetic development, I moved to another synthetic biology subfield that heavily relies on SJRs: synthetic immunotherapy ^2, 15–18, 21, 35, 36^. Due to their high spatial control, modularity, and recent humanization, SJRs are rapidly accelerating towards clinical use ^2, 15–18, 21, 35, 36^. As a result, SJR-based direct activation circuits currently dominate the subfield, enabling programming T cells with novel behaviors such as multi-antigen discrimination (logic gating) ^14–16, 21, 36^. Such behavior in T cells has powerful implications; the tumors targetable can be broadly expanded as there is no longer the need to rely on a single specific antigen ^2, 3, 11, 12, 15–17, 21, 40^. These logic-gated synthetic biology (SB) CAR T cells have been deployed *in vivo* with a prime and kill strategy against two different types of tumors with either heterogenous or homogenous antigen expression ^16, 21, 36^.

In the prime and kill strategy against a heterogenous tumor, SB CAR T cells have an SJR that, in response to tumor antigen one, induces expression of a chimeric antigen receptor (CAR) against tumor antigen two. This direct activation circuit has highly localized anti-tumor effects; SB CAR T cells primed by dual antigen positive tumor cells killed both dual antigen and neighboring single antigen positive tumor cells ^21, 36^. Targeting a homogenous tumor is similar, but SB CAR T cells are primed by antigen on proximal healthy cells instead ^21^. In both cases, killing was spatially restricted to the tumor area and SB CAR T cells required both antigens present to mediate tumor clearance.

Against either type of tumor, the directly activated SB CAR T cells are highly dependent on the spatial presence of tumor antigen one/priming antigen to maintain CAR expression and kill tumor cells ^21, 36^. I therefore tested if CAR amplification and prolonged duration of expression conferred by the SJR amplifier circuits could improve tumor killing ^16^. As the direct activation circuit in SB CAR T cells is effectively the direct activation circuit in the 3-layered structure and multipole structure (Fig.3-4), but with the CAR replacing the cadherin/blue ligand as the target gene, I could therefore modify the directly activated gray cells from those experiments, along with the amplifier imbued cells, to create a simple but highly modular setup for testing these cells against these tumors.

To target a heterogenous tumor with the prime and kill strategy, I reprogrammed the cadherin expressing gray cells into SB CAR T cells by coding the SJR to respond to yellow tumor antigen one and replacing the cadherin/blue ligand gene with anti-(red tumor antigen two) CAR (*ARA CAR*) (Fig.6A).

Yellow tumor antigen one represents tumor antigen one and red tumor antigen two represents tumor antigen two (Fig.6C). In directly activated gray SB CAR T cells, the SJR directly drives CAR expression (Fig.6A) whereas in the amplifier imbued SB CAR T cells (Fig.6B), the SJR drives the amplifier which then drives CAR expression. I then created the remaining cells with the appropriate antigen combination. Surrounding (S) cells have neither tumor antigen one (yellow antigen) nor tumor antigen two (red antigen) on their surface and they represent normal healthy cells around and within the tumor (Fig.6C). Dual antigen (DA) tumor cells have yellow tumor antigen one and red tumor antigen two on their surface (Fig.6C). Single antigen tumor cells (U) have the red tumor antigen (Fig.6C). Tumor cells can receive pro-apoptotic signals from SB CAR T cell CAR signaling as is *in vitro* ^41–43^.

Then, by seeding different ratios of these four cell types, I could create a variety of heterogenous tumors (Fig.6C). Initial testing with two cell ratios differing primarily in SB CAR T cell number yielded similar results (SFig.7). Temporarily amplified activated (TAA) and permanently amplified activated (PAA) SB CAR T cells had more CAR^+^ cells than the directly activated (D) cells (SFig.7A). Confirming the amplification and temporal behavior of these circuits, more gray cells became CAR^+^ and maintained being CAR^+^ over time (SFig.7B). The direct activation circuit, as expected, had the majority of gray cells become CAR^+^ but they reverted to CAR^-^ as time passed (SFig.7B). Cells with either amplifier appeared to kill more tumor cells than the directly activated SB CAR T cells (SFig.7C). I validated that these SB CAR T cells required, as is observed *in vitro* and *in vivo* ^21^, both antigens to kill tumors cells (SFig.9).

**Figure 7.**
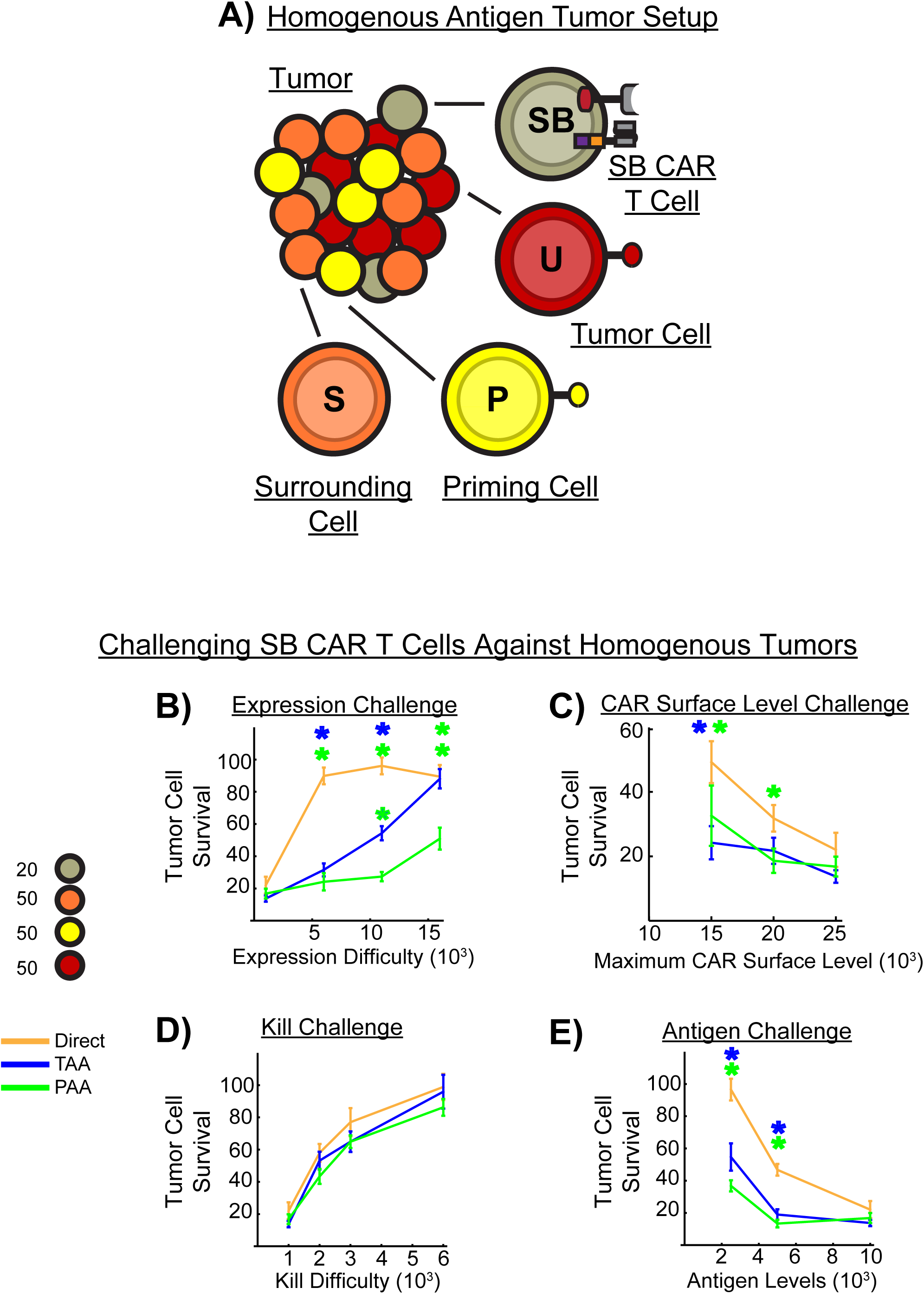
SJR Amplifier Circuits Improve CAR T Cell Killing Against Homogenous Tumors Across Several Conditions. A) Creation of the homogenous antigen tumor setup. The SJR direct activation circuit from Fig.6A was tested against a homogenous tumor *in vitro* ^21^ thus I modified the heterogenous tumor in Fig.6C to homogenous by converting pink dual antigen DA tumor cells to yellow priming P healthy cells. Other cells (S, U, SB CAR T) remained the same. Mixing these cells enables creating a variety of homogenous tumors. Tumor here is 19.8±0.85 gray SB CAR T cells with 55.7±2.13 orange S cells with 52.5±2.33 yellow P cells and 51±1.82 red U cells. Circuit and curve color key are given to the left of the bottom plots. B) CAR Expression Challenge. SB CAR T cells were challenged with increasing difficulty of CAR expression. In general, using an amplifier to amplify and prolong CAR expression decreases tumor cell survival more than the direct activation circuit does. C) CAR Surface Level Challenge. SB CAR T cells were challenged with different maximum saturation CAR surface levels. At lower maximum CAR surface levels, amplifier imbued SB CAR T cells decreased tumor cell survival more than the directly activated cells did. D) Kill Challenge. SB CAR were challenged to kill increasingly CAR signaling resistant tumor cells and all circuits had similar levels of tumor cell killing, indicating that amplifiers do not confer a tumor killing advantage in this homogenous antigen tumor setup. E) Antigen Challenge. SB CAR T cells were challenged to kill tumor cells with less priming antigen available. Amplifier imbued SB CAR T cells could amplify the lower signal to drive CAR expression and kill more tumor cells. N=10 for each circuit for each parameter tested. * denotes significance and is colored the curve it differs from. Simulations run for 50,000 timesteps with shown data from this endpoint.

These results indicate that amplifier imbued SB CAR T cells can improve heterogeneous tumor killing compared to the directly activated cells, but this advantage is moderate. The tumor survival difference was ∼10% and overall, all circuits had potent tumor killing (SFig.7C). In these trial experiments, I had used the optimal setup for SB CAR T cell performance: CAR expression was very easy (expression difficulty, β CAR=1000), CAR surface saturation levels high (maximum CAR surface level, κ=25000), tumor cells were very easy to kill (pro-apoptotic protein expression difficulty as a result of CAR signaling, β apoptosis=1000), and priming antigen levels high on the corresponding cells (L=10000). To determine where the amplifier circuits would confer a potent advantage to SB CAR T cell tumor killing, I subjected the SB CAR T cells to a gauntlet of challenges based on these parameters (Fig.6D-G).

A common challenge in engineering synthetic circuits is sufficient expression of the induced target gene, often requiring several iterations of the construct to achieve the desired behavior ^44–46^. I therefore challenged these SB CAR T cells by increasing the expression difficulty of the CAR via increasing the parameter β CAR (Expression Challenge, Fig.6D). At the easiest expression difficulty (β CAR=1000), both the temporarily amplified activated (TAA) and directly activated (D) SB CAR T cells had similar performance, but the latter quickly lost tumor killing capabilities as CAR expression difficulty increased (Fig.6D). Temporarily amplified activated SB CAR T cells decreased tumor cell survival more than the directly activated cells did at higher difficulties (Fig.6D). Permanently amplified activated (PAA) SB CAR T cells managed to maintain superior tumor killing over the other circuits as expression difficulty increased (Fig.6D). These results indicate that when targeting a heterogenous tumor, if CAR expression difficulty is a concern, using an amplifier to amplify and prolong CAR expression could improve tumor killing. Furthermore, using the permanently amplified activation circuit could be additionally advantageous to using the temporarily amplified activation circuit.

Alternatively, CAR T cell anti-tumor capabilities can be impaired by insufficient surface levels of CARs ^47–49^. To determine how this affects amplifier imbued SB CAR T cells, I challenged the cells by testing different CAR surface saturation levels, modeled by parameter κ in GJSM (CAR Surface Level Challenge, Fig.6E). Expression of CAR was kept easiest (β CAR=1000) and overall, both amplifier circuits outperformed the direct activation circuit (Fig.6E). These results indicate that against a heterogenous tumor, when CAR surface levels saturate low, amplifier circuits can yield superior tumor cell killing.

The two above challenges stem from SB CAR T cell engineering, but the tumor cells themselves can hinder killing ^43, 50–57^. Tumor cells can resist CAR T cell mediated apoptosis via alteration to the death receptor pathways ^41–43^. This can be modelled in my setup by increasing the parameter β apoptosis, which models pro-apoptotic protein expression as a result of CAR signaling (Kill Challenge, Fig.6F). The amplifier circuits overall appeared to decrease tumor cell survival more than the direct activation circuit, but the difference was moderate, ranging from 10%-20% (Fig.6F). As kill difficulty increased, tumor cell survival increased as expected (Fig.6F). These data indicate that when tumor cells become more resistant to CAR T cell cytotoxicity, amplifying and prolonging CAR expression could confer a moderate advantage.

In an alternative but common mechanism, tumor cells can evade CAR T cells by downregulating tumor antigen expression ^17, 51–53, 55, 56^. I therefore tested the SB CAR T cells against tumors with varying levels of yellow tumor antigen one. At the lowest antigen level (L=2500), amplifier imbued SB CAR T cells decreased tumor cell survival more than the directly activated cells did, with the permanently amplified activated cells further outperforming the temporarily amplified activated cells (Fig.6G). As the antigen level increased, performance difference between all three diminished but permanently amplified activated cells maintained significantly decreased tumor survival compared to the directly activated cells (Fig.6G). Thus, amplifying and prolonging CAR expression can improve tumor killing where there is tumor antigen downregulation. At low antigen levels, the permanently amplified activation circuit can yield superior tumor killing compared to the other two circuits.

### SJR amplifier circuits improve CAR T cell tumor killing against homogenous tumors

To target a homogenous tumor with the prime and kill strategy, healthy tissue antigen is used to prime rather than a tumor antigen. SB CAR T cells remain programmed with the direct activation circuit, but the SJR now responds to yellow priming antigen to express a CAR against the tumor antigen. Then, these SB CAR T cells can be primed by a healthy cell’s yellow antigen to kill nearby tumor cells expressing the tumor antigen ^21^.

In my setup, this can be modeled by simply shifting the antigen expression in the heterogenous tumor. I altered dual antigen (DA) tumor cells to become priming (P) cells that only express the yellow priming antigen (Fig.7A). These cells represent healthy cells that serve to prime the SB CAR T cells (Fig.7A). SB CAR T cells remained the same. Directly activated SB CAR T cells retain the SJR that responds to yellow antigen and induces expression of anti-(red antigen) CAR (*ARA CAR*) (Fig.6A). Amplifier imbued SB CAR T cells retain the anti-(yellow antigen) SJR that drives the amplifier rather than the target gene (Fig.6B). Surrounding (S) cells and tumor (U) cells represent tumor proximal, non-antigen expressing healthy cells and single antigen (red antigen) tumor cells, respectively (Fig.7A).

Like in the heterogenous tumor setup, I could seed different ratios of these four cell types to create a variety of homogenous tumors. I performed the same tests as those against the heterogenous tumor. Both the temporarily amplified activation (TAA) and permanently amplified activation (PAA) circuits were able to yield more CAR^+^ cells than the direct activation circuit (D) (SFig.8A). However, the difference was small (∼5%) (SFig.8A). Tumor cell survival appeared to differ between circuits; SB CAR T cells with either the temporarily amplified activation circuit or permanently amplified activation circuit appeared to reduce tumor cell survival more than cells with the direct activation circuit did (SFig.8C).

**Figure 8.**
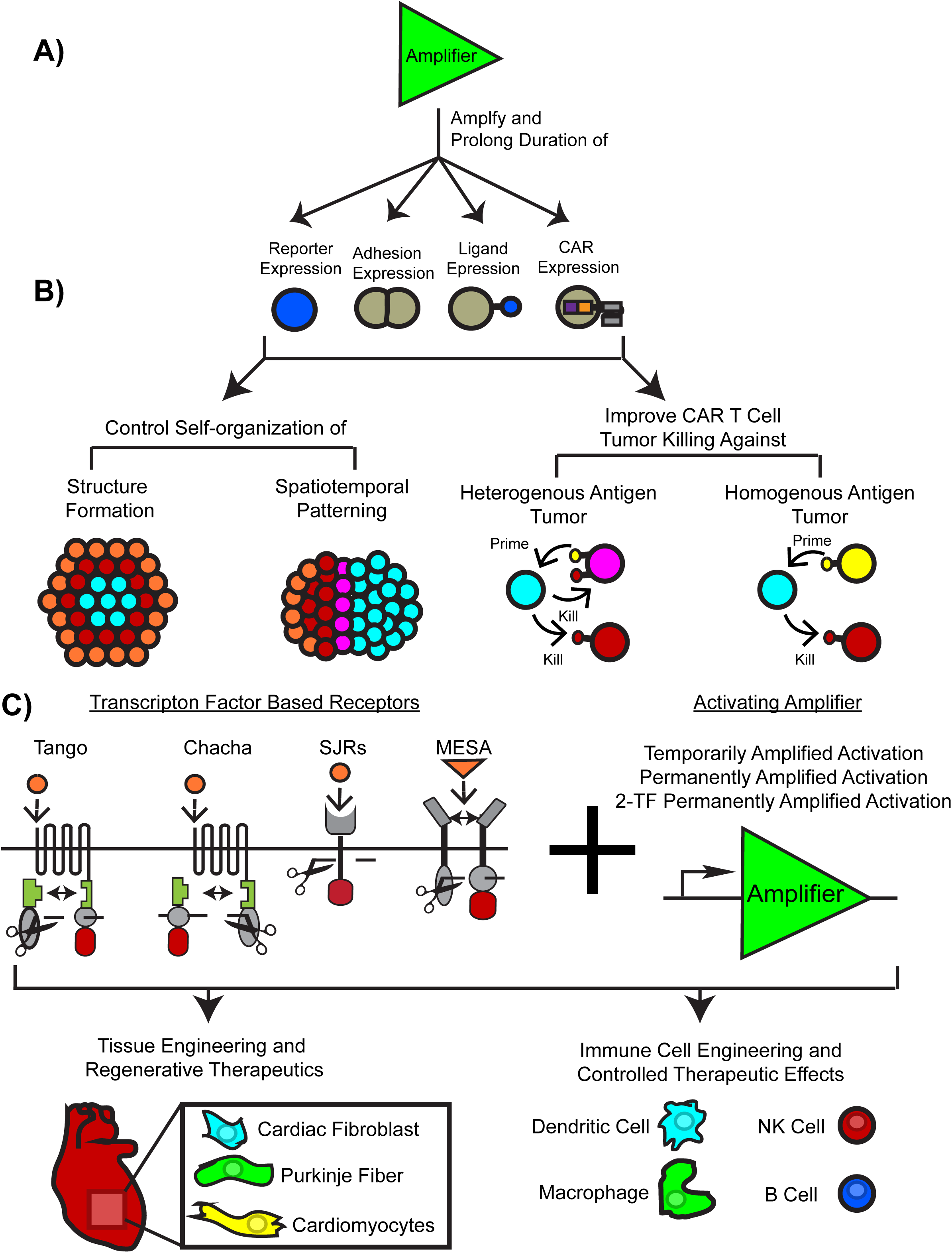
Amplifiers Expand Control in Synthetic Circuits for Self-organization and Immunotherapy. A) I demonstrate here that amplifiers enable amplification and temporal regulation of target gene expression, thus expanding the control of synthetic circuits. B) When these amplifiers are combined with SJRs and control genes for self-organization or immunotherapy, they improve engineered self-organization, modularly direct spatiotemporal patterning, and improve CAR T cell killing against heterogenous and homogenous antigen expressing tumors. C) The experiments here focus on SJRs, but the amplifiers should be compatible with other transcription factor based synthetic receptors. These synthetic receptors and amplifier based circuits, in principle, broadly expand control in synthetic development efforts, furthering understanding of self-organization and therapeutic efforts in tissue engineering and regenerative medicine as well. These circuits should be also applicable to immunotherapy, potentially broadening control over immune cell behavior for synthetic immunotherapy.

However, the percentage difference was ∼5% nor were results significant in the tumor with less SB CAR T cells (SFig.8C).

These results are similar to those obtained in the initial tests with the heterogenous tumor, and again suggest that the amplified SB CAR T cells can improve tumor killing, but likely have a moderate advantage over directly activated cells. This test configuration kept the optimal parameters for SB CAR T cell performance: CAR expression difficulty easiest, β CAR=1000, CAR surface saturation level high, κ=25000, pro-apoptotic protein expression difficulty as a result of CAR signaling easiest, β apoptosis=1000, and priming antigen level high (L=10000). I therefore subjected these SB CAR T cells to the same battery of challenges performed in the heterogenous tumor setup (Fig.7B-E).

When challenged with increased difficulty of CAR expression, all activation circuits had similar performance at the easiest CAR expression difficulty (β CAR=1000), but directly activated (D) SB CAR T cells quickly decreased in tumor killing as CAR expression difficulty increased (Fig.7B). SB CAR T cells with the temporarily amplified activation circuit (TAA) decreased tumor cell survival more than the direct activation circuit did at intermediate difficulties (β CAR=6000 and 11000) but faltered at the highest difficulty tested (β CAR=16000) (Fig.7B). In contrast, permanently amplified (PAA) SB CAR T cells maintained superior tumor killing as difficulty increased (Fig.7B). These results indicate that in the prime and kill strategy against a homogenous tumor, using an amplifier to amplify and prolong CAR expression can overcome challenging CAR expression and decrease tumor survival. Moreover, where CAR expression is especially challenging, using the permanently amplified activation circuit can improve tumor clearance.

Pitting the SB CAR T cells in the CAR Surface Level Challenge (Fig.7C) revealed that amplifier imbued cells, at the lowest surface saturation level (κ =15000), significantly outperformed the directly activated cells (Fig.7C). At the intermediate difficulty (κ =20000), amplified activated SB CAR T cells performed similarly to directly activated SB CAR T cells, while permanently amplified activated SB CAR T cells decreased tumor survival more than directly activated cells did (Fig.7C). Therefore, in this healthy tissue priming strategy, where CAR surface levels saturate low, using amplifiers to amplify CAR expression can decrease tumor survival.

I then tested these SB CAR T cells against CAR T resistant tumor cells with the Kill Challenge (Fig.7D). At all difficulties tested, all SB CAR T cells performed similarly regardless of the activation circuit type (Fig.7D). No significant difference was detected between any circuits at the same kill difficulty level. As kill difficulty increased, tumor cell survival increased as expected (Fig.7D). These data indicate that in the prime and kill strategy against a homogenous tumor, unlike against a heterogenous tumor (Fig.6F), using an amplifier will not confer a tumor killing advantage.

Finally, I tested the SB CAR T cells in the Antigen Challenge (Fig.7E). At the lower priming antigen levels (L=2500 and L=5000), amplifier imbued SB CAR T cells decreased tumor cell survival more than the directly activated cells did (Fig.7E). Neither the temporarily amplified activated nor permanently amplified activated SB CAR T cells outperformed one another and performance difference between all circuits diminished as antigen levels increased (Fig.7E). Thus, where priming antigen is low, amplifying and prolonging CAR expression with either amplifier circuit could improve tumor cell killing.

## DISCUSSION

Amplifiers that amplify target gene expression and enable unidirectional temporal regulation expand the control of the circuits commonly used in synthetic development and synthetic immunotherapy. Here I show via mathematical and *in silico* analysis that the activating amplifiers amplify target gene expression and enable prolonging duration of target gene expression, with different amplifiers enabling differing levels of temporal control (Fig.8A). In synthetic development, amplifiers combined with SJRs improve target structure formation and enable modular spatiotemporal control (Fig.8B). In synthetic immunotherapy, these amplifier circuits can improve synthetic biology (SB) CAR T cell tumor killing against both heterogenous and homogenous antigen expressing tumors (Fig.8B). Together, these results demonstrate the capabilities and potential use of amplifiers in not just basic applications, but also clinical applications as well (Fig.8C).

To study the capabilities and applications of these SJR-amplifier circuits, I used the GJSM model implemented in a cellular Potts model ^26, 33, 34^. This model combination has been previously validated for predicting self-organization and patterning driven by SJRs ^33^. The two synthetic development structures (3-layered and multipole) that have been further examined in this study were originally modelled *in silico* by this combination ^33^. Although this framework was not previously validated for synthetic immunotherapy, the setup I created to model SB CAR T cells and the tumors yielded results surprisingly similar to those obtained biologically ^21, 36^. Directly activated *in silico* SB CAR T cells strongly killed tumor cells (SFig.7-8) as is observed *in vitro* ^21^. These *in silico* cells were highly dependent on SJR signaling to become CAR^+^ (SFig.9) and maintain being CAR^+^ (killing of DA cells resulted in reversion to CAR^-^, SFig.7). These are all features observed *in vitro* ^21^. Therefore, while numerous factors remain to be included and refined (i.e. cytokine secretion and signaling, T cell activation mediated expansion, exhaustion, etc), this simple setup provides a viable platform for computationally testing a variety of SB CAR T cells against a variety of tumors. Developing this platform could expand *in silico* methods in synthetic immunotherapy and create a pipeline similar to how GJSM is currently used to test circuit designs for synthetic development ^33^. While computational efforts for synthetic immunotherapy begin with this study, similar efforts are at least underway for regular CAR T cell immunotherapy ^58–62^.

An additional advantage of using a mathematical and *in silico* approach is that it enabled me to rapidly yet thoroughly investigate the amplifiers and their circuits. Changing parameters (i.e. type of gene expressed, expression difficulty, surface saturation levels) was simply changing code while reengineering cells *in vitro* would likely take years with one researcher as in this study. However, using a computational approach does give a theoretical analysis of the circuits. For example, transcription factor feedback loops similar to those in the permanently amplified activation circuit (Fig.1F and 6B) do not always maintain permanent gene expression *in vitro* ^32, 63–66^. Some cells can lose permanent gene expression over the course of multiple generations, and this is hypothesized to be a combination of epigenetic silencing and/or failure of daughter cells receiving sufficient transcription factors post mitotic division ^32^. Depending on the purpose of the circuit, this feature may be desirable. Permanently amplified activated SB CAR T cells increase tumor cell killing but theoretically can increase the risk of on-target off-tissue toxicity. Loss of CAR expression can mitigate this and continued expression of the SJR allows these cells to spatially reactivate where they are again exposed to the tumor. If undesirable, this limitation could be overcome by increasing transcription factor levels before division ^32, 64^ or reactivating silenced genes through another synthetic circuit ^67, 68^.

In this study, I designed the circuits generically (i.e. ATF1, anti-yellow antigen SJR, etc) such that there is no specific SJR (i.e. anti-CD19 synNotch) or specific transcription factor (i.e. LexA-VP64) modelled. I have deliberately left it to the user to choose the specific components for the circuits. For example, the specialized doxycycline/hypoxia/DNA damage based circuits designed by Burrill et al used a trigger circuit with zinc finger transcription factors to drive their feedback loop ^32^. Keeping the amplifier circuits generic enables their compatibility with future tools for as synthetic receptors ^17^ and transcription factors ^69^ are continually expanded, reengineered, and improved, the breadth of components for the amplifier and thus amplifier circuits will expand as well.

Circuit generalizability should additionally extend amplifier compatibility beyond SJRs to other transcription factor based receptors as well. As the amplifiers are transcription factor based, they should be compatible with other transcription factor based receptors such as modular extracellular sensor architecture (MESA) and Tango/ChaCha ^70–78^ (Fig.8C). MESA is a class of fully modular synthetic receptors that dimerize in response a choice soluble ligand to release a choice transcription factor ^72, 74, 75, 78^ while Tango/ChaCha are modified GPCRs that bind a soluble ligand to release a choice transcription factor ^73, 77^. Then, having these receptors release a transcription factor that drives the amplifier, rather than the target gene, should enable amplification and prolongation of target gene expression as is observed with SJRs. In theory, these amplifiers should be broadly compatible with transcription factor based circuits overall (Fig.8C).

The amplifiers and SJR amplifier circuits designed in this study serve as generic yet flexible tools for improving control in synthetic biology. The ability to add gene amplification and prolong duration of gene expression expands the capability of SJR based circuits and likely other transcription factor based circuits as well. Though the experiments here focused on adhesion-based self-organization and T cell immunotherapy, combining these amplifiers with different synthetic receptors, in principle, broadly expands control over processes such as tissue regeneration ^19^ or other immune cell behavior ^79, 80^ (Fig.8C). In the upcoming accompanying paper, I demonstrate the ability of these amplifiers to drive other amplifiers and obtain inhibitory gene expression that shortens or turns off duration of gene expression.

## ACKNOWLEDGEMENTS

I would like to thank UNMC’s library services for access to research articles and UNMC’s printers for allowing me to print articles.

## AUTHOR CONTRIBUTIONS

C.L. conceived, designed, performed, analyzed, and wrote the entire study.

## METHODS

### Key Resource Table

| REAGENT or RESOURCE | SOURCE | IDENTIFIER |
| --- | --- | --- |
| Software and Algorithms |  |  |
| CompuCell3D (CC3D) v3.7.8 | <sup>34</sup> | RRID:SCR_003052 |
| Mathematica v12.0.0.0 | Wolfram Research | RRID:SCR_014448 |
| JMP v11.0.0 | SAS Institute | RRID:SCR_014242 |
| Excel v2301 | Microsoft | RRID:SCR_016137 |
| General Juxtacrine Signaling Model (GJSM) in CC3D | <sup>33</sup> | N/A |

### Lead Contact

Requests for information, resources, and code will be fulfilled by Calvin Lam (<u>calvin.lam.k@gmail</u>).

### The *in silico* cell line ISL929 in CompuCell3D

ISL929 is an *in silico* cell line developed in the cellular Potts modeling software CompuCell3D (CC3D) and has been previously combined with the Generalized Juxtacrine Signaling Model (GJSM) to successfully predict synthetic juxtacrine receptor (SJR) driven self-organization ^33, 34^. As ISL929 is specifically designed to model the *in vitro* mouse fibroblast cell line L929 used in the reference biological self-organization experiments (3-layered and multipole) ^9^, I chose to use ISL929 as my *in silico* cell line. I provide a brief description of ISL929 below and the original *in silico* study that developed this line is given here ^33^.

ISL929 is implemented in CC3D as 3D multipixel cells that have physical properties such as surface area, volume, adhesion, and motility. ISL929 cells are programmed to desire a spherical morphology as is observed in suspension *in vitro* ^9, 33^. Each cell (σ) is assigned a desired radius (DR) drawn from a Gaussian distribution with mean of 3 pixels and standard deviation 0.5 pixels. The desired surface area (DS) and desired volume (DV) are then calculated from this desired radius (4πr^2^ for surface area and 4/3πr^3^ for volume), promoting cells to want a spherical morphology. How strongly the cell desires this spherical morphology is determined by the parameters λ_S_ for the desired surface area and λ_V_ for the desired volume. These parameters can be thought of as the spring constants in Newtonian physics. Then, a population of ISL929 is relatively homogenous in morphology, overall consisting of spherical cells but with slight size differences due to differing in desired radius. For ISL929, λ_S_ and λ_V_ are set to 2.2, generating cells that are spherical as is observed with *in vitro* L929 cells ^9, 33^. However, if ISL929 cells are modified with different circuits that change adhesion, these parameters can change as determined in both the *in vitro* and *in silico* study ^9, 33^. Full parameters are given in Supplementary Table 1.

With the basic morphology of ISL929 cells defined, growth and division can be implemented.

Growth is achieved by subjecting the desired radius DR to fluctuations drawn from a uniform distribution, with a slight skew towards increasing the desired radius (-3*10^-3.88^ to 4*10^-3.88^). Cells therefore slowly increase their desired radius, increasing the desired surface area and volume as well. When cells reach the threshold volume (2*4/3πr^3^ with r=3 pixels), cells undergo symmetric division into two cells. After division, the parent cell is assigned a new desired radius and undergoes growth once again. The child cell is assigned the same parameters. With these rules and parameters, ISL929 roughly doubles every 24000 timesteps and as *in vitro* L929 cells roughly double every 24 hours, this sets 24000 simulation timesteps to 24 hours in real time ^9, 33^.

*In vitro* L929 cells weakly adhere to one another and in CC3D, cell-cell adhesion is defined by a matrix of parameters J. Cell adhesion to different cells (i.e have cadherin or different cadherin expression) have different J values. However, as *in vitro* L929 weakly adhere and do not express any cadherin, ISL929 have basal adhesion value J=49 to any other cell type, cadherin expressing or not. This is defined relative to the media such that J=49 sets ISL929 cells to weakly aggregate to one another in the simulated media. Like with λ_S_ and λ_V_, however, these J values can change as ISL929 cells are modified with different circuits that drive different cadherins ^9, 33^. Full parameters are given in Supplementary Table 1.

Cell motility in CC3D is defined by parameter T and as strong adhesion (i.e. cadherin expression) is generally abstracted to decrease cell motility ^81–83^, ISL929 motility is defined by the formula

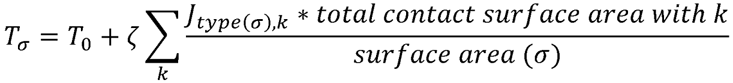

Each cell σ is assigned its own T value according to this formula, with T_0_ being basal motility and ζ being the weight of adhesion attenuating motility. The summation term determines the motility of the cell due to the adhesivity of its local environment. K designates the types of cells to sum over, with types of cells stratified by surface expression of cadherins (i.e. one type of cell is high E-cadherin^+^ while another could be low E-cadherin^+^). For example, a simulation with high E-cadherin^+^ ISL929 and low E-cadherin^+^ ISL929 would have two cells types and thus k sums up to two. J_type(σ),k_ is the adhesion value of the type of focal cell σ to cells of type k. For example, if the focal cell is of parental ISL929 type, then its J value is 49 to any other cell type k. This J value is then fractionalized to the total contact surface area with cells of type k by dividing over the focal cell’s surface area (surface area (σ)). Higher contact area with less adhesive cell types (higher J value) results in a higher value from the summation term, yielding higher motility. Maximum motility is obtained in the medium, with J=52 for cell to medium adhesion. Overall, this motility formula allows a cell to sense the adhesivity of its local environment and attenuate its motility accordingly. In general, more adhesive environments lower motility while less adhesive environments restore motility. Parental ISL929 have T_0_ set to 100, ζ set to 0.5, and J set to 49, resulting in highly motile cells as is observed *in vitro* ^9, 33, 84^. Full parameters are given in Supplementary Table 1.

With the physical and basic properties of ISL929 defined (all the properties described thus far are identical to those in the original computational study ^33^), cell motion can finally be described. Cells move in the cellular Potts model by “pixel copy attempts”. That is, cells attempt to move by copying their pixels over to a neighboring pixel ^26, 33, 34^. The success probability of this move is determined by the probability function P=e^-ΔH/T^, where P is the probability of success, ΔH the change in global system energy calculated from all the pixel copy attempts at said timestep, and T the focal cell motility. As ISL929 has surface area, volume, and adhesion, H takes the form of

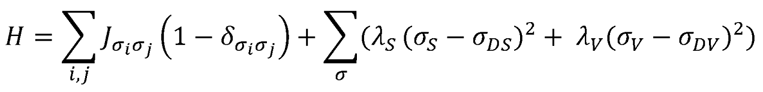

as in ^26, 33^. σi and σj are the identities of cells that occupy pixel site i and j, respectively. J is the adhesion value of the types of cells of σ_i_ and σ_j_. The Kronecker delta (δ) term limits the calculation to cell-cell interfaces. λ_S_ and λ_V_ controls the cell’s desire to achieve its desired surface area (σ_DS_) and desired volume (σ_DV_), respectively. σ_S_and σ_V_ are the cell’s actual surface area and volume, respectively, at a given timestep.

### Brief overview of Generalized Juxtacrine Signaling Model (GJSM)

Generalized Juxtacrine Signaling Model (GJSM) is a mathematical model developed and validated to describe synthetic juxtacrine receptor (SJR) regulation of a target gene’s expression ^33^. SJR activation of target gene expression is described by the equation

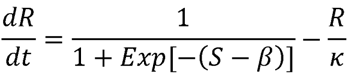

The change in target gene protein level (R) at a given timestep is a balance between production from SJR signaling (first term) and loss from degradation (second term). In the first term, production is calculated from signal (S) and target gene expression difficulty (β). Because SJRs and transcription factors activate expression in this study, S models SJR or transcription factor signaling that drive the amplifier and/or target gene. However, as it is suggested that synthetic transcription factor based circuits are bistable (either sufficient or insufficient transcription factor to drive expression ^32, 64^), S should be weighed against an expression difficulty parameter β. β models the difficulty of gene expression, encompassing a variety of biological factors such as promoter, transcription, and/or translation inefficiency ^33^. The logistic form of the production term (Exp is e) smoothly limits production from 0 to 1 per timestep, representing minimal to maximal production at a given timestep.

The degradation term is the standard linear decay rate commonly deployed in biochemistry models. Target gene protein level R decays proportionally to itself and inversely to decay constant κ. κ controls the protein saturation level. For example, in the synthetic development experiments (3-layered and multipole, Fig.3-5), it can control the maximum level of cadherin and/or blue ligand while in the synthetic immunotherapy experiments (Fig.6-7), it can control maximum CAR surface level. Then, the degradation term also ranges from 0 to 1 per timestep, representing minimal to maximal degradation at a given timestep.

### The SJR amplifier circuits as modelled by GJSM

With the base equation described, the equations for the activation circuits (direct activation circuit, temporarily amplified activation circuit, permanently amplified activation circuit, and 2-TF permanently amplified activation circuit) can now be defined.

#### Direct Activation Circuit

Because the direct activation circuit is the SJR driving target gene expression directly (Fig.1D), the circuit equation is

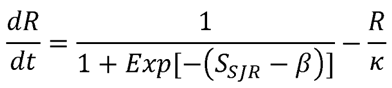

This form is identical to those in previous direct activation circuits ^33^ and parameters are as described in the above section. S_SJR_ is the number of activated SJRs (binding cognate ligand/antigen to release its transcription factor) at a given timestep. See how S_SJR_ is calculated in the below section, Adding ligands, receptors, and the activation circuits into ISL929.

#### Temporarily Amplified Activation Circuit

In the temporarily amplified activation circuit, the SJR drives the expression of an activating transcription factor (ATF) that then drives expression of the target gene (Fig.1E). Because this is a two-step process, two equations are required. First, ATF expression driven by the SJR is described as following.

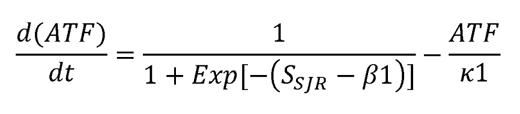

ATF production is driven by SJR signal S_SJR_. β1 controls the expression difficulty for the SJR to drive ATF expression while κ1 controls the maximum/saturation level of ATF protein.

As the ATF drives the expression of the target gene, target gene product level (R) is described by the equation S_ATF_ is the ATF levels (ATF from the above equation) at a given timestep and drives the expression of the target gene. β2 controls the expression difficulty for the ATF to drive target gene expression while κ2 controls the maximum/saturation level of target gene protein.

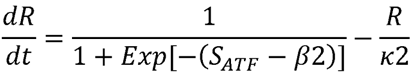

#### Permanently Amplified Activation Circuit

In the permanently amplified activation circuit, the SJR drives the expression of an activating transcription factor (ATF) and the target gene (R). This ATF also drives expression of itself and thus also the target gene R (Fig.1F). The equations are as following.

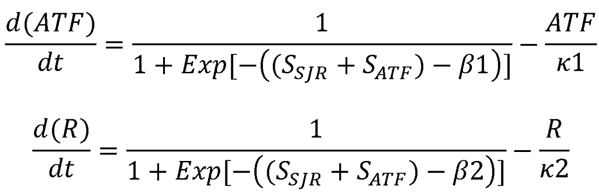

S_ATF_ is the ATF levels (ATF) from solving the first equation.

#### 2-Transcription Factor (2-TF) Permanently Amplified Activation Circuit

In the 2-transcription factor (2-TF) permanently amplified activation circuit, the SJR drives the expression of two activating transcription factors, ATF1 and ATF2, on the same gene cassette. Then, ATF1 and ATF2 are assumed stoichiometrically equal and have the same equation.

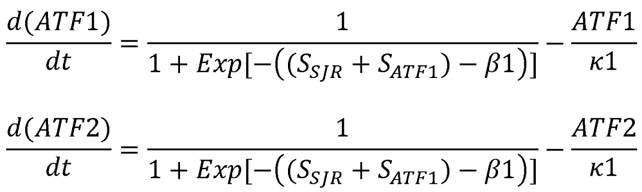

The parameters of ATF1 and ATF2 (β1 and κ1) are kept the same in this study to decrease number of parameters used but can be changed depending on the transcription factors modelled or inefficiencies of the transgene cassette (i.e. IRES inefficiency). Target gene product levels R are driven only by ATF2 and thus are described by the equation.

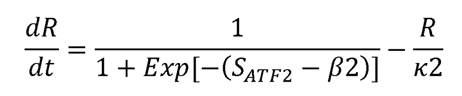

The parameters tested in this study for all the above equations are given in Supplementary Table 1. It is important to note that within a set of equations representing an activation circuit, the parameters β and κ can differ between equations due to differences in expression difficulty and degradation.

### Adding ligands, receptors, and the activation circuits into ISL929

With ISL929 cells defined and the mathematical equation of the activation circuits described, I could finally implement the equations into the cells to generate the circuit imbued ISL929 for the synthetic development and synthetic immunotherapy experiments (SFig.1A). I then added constitutive ligand expression onto the appropriate cells (i.e. orange ligand expression on orange cells, red antigen on single antigen positive tumor cells). I simplified the time dependent equation from the original *in silico* study ^33^ and set it to a constant value L on the cell’s surface per simulation. Constitutive SJR expression was then added to the appropriate cells (i.e. anti-(blue ligand) SJR on orange cells, anti-(yellow antigen) SJR on gray SB CAR T cells). As there is no evidence of SJR to be limiting in the reference experiments, SJR surface levels was assumed to be in excess and thus not needed for the subsequent calculations ^33^. However, if desired, the full formulation of GJSM can easily model this ^33^. Parameters for ligand/antigen levels per experiment are given in Supplementary Table 1.

Having defined the constitutive ligand/antigen levels, it becomes possible to calculate the signal from SJR signaling, S_SJR_. In the most generic form, SJR signaling is dependent on both the ligand/antigen level the focal cell is exposed to and the SJR receptors on the focal cell that can respond to these ligands/antigens. In this study, because SJR levels are assumed to be non-limiting and because SJRs function in a 1:1 stoichiometry with its ligand (one SJR binds one ligand to release one transcription factor) ^16, 17, 19, 33^, SJR signal S_SJR_ depends only on the amount of ligand/antigen the focal cell is exposed to (SFig.1B). This can be calculated by the equation below.

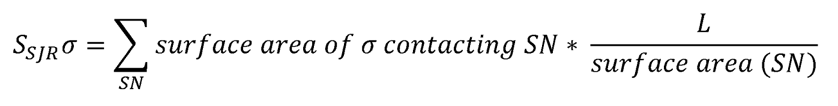

At a given timestep, focal cell σ receives SJR signal S_SJR_σ calculated from the total cognate ligand/antigen level it is exposed to. Signaling neighbor (SN) is a neighboring cell expressing the appropriate cognate ligand (i.e. orange ligand on orange cells cognate to anti-(orange ligand) SJR on a focal gray cell). For a signaling neighbor SN, the amount of cognate ligand/antigen it presents is calculated by multiplying the surface area the focal cell σ is in contact with the signaling neighbor by the ligand density on said signaling neighbor (L/surface area(SN)). For a constitutive ligand, L is defined as a constant value while for a ligand that is a target gene (i.e. blue ligand), L is defined by the value of R. Repeating this calculation over each signaling neighbor sums the cognate ligand/antigen level the focal cell is exposed to at a given timestep and gives S_SJR_σ at that timestep. While not required in this study, cells with multiple SJRs will have several S_SJR_σ calculated depending on the cognate neighbors present at a given timestep.

### Linking target gene expression to behavior

To link target gene expression to the intended behavior (i.e. cadherin expression, ligand expression, CAR expression), I used a discrete state-transition model as in the original study and commonly deployed in computational biology ^26, 33, 85, 86^. If target gene protein level R exceeds a threshold (7000 for all simulations here), then the cell gained the feature of the target gene. Cells with blue ligand level that exceeded the threshold became blue and could signal to cells with anti-(blue ligand) SJR. Cells with high E-cadherin levels that exceeded the threshold became highly adhesive (J changed). Cells with CAR levels that exceeded the threshold could target the cognate tumor cells, a bistability feature that is observed *in vitro* ^47–49^. Tumor cells, upon accumulating enough pro-apoptotic proteins as a result of SB CAR T cell signaling, irreversibly commit to apoptosis ^43, 87–92^. Where cells did not irreversibly fate commit (i.e. cadherin expression, ligand expression, CAR expression), falling under this threshold reverted cells, losing the behavior. Full parameters are given in Supplementary Table 1.

An example schematic of the entire signaling process, from constitutive ligand expression to SJR signaling to amplifier expression to target gene expression to resulting behavior change, is given in SFig.1B.

### General simulation conditions

At the start of the simulation, cells were initialized as a 5×5×5 pixel cube in a 100×100×100 pixel lattice. Initial configuration and ratio of cells is specified in the appropriate section below. Data was collected every 100 timesteps for analysis.

### Homogeneity Index

To quantify the quality of the 3-layered structure’s core and the quality of the multipole structure’s poles, I used the homogeneity index defined in ^33^. This measure was previously used to quantify self-organization in the simulated reference structures ^33^. As the core forms from high E-cadherin^+^ cells (blue/cyan) and the poles form from either N-cadherin^+^ cells (blue/cyan) or P-cadherin^+^ cells (red/pink), I was interested in the contact between cadherin cells to other same cadherin cells. For example, in the 3-layered structure, I was interested in high E-cadherin^+^ cells contacting other high E-cadherin^+^ cells while in the multipole structures, I was interested in N-cadherin^+^ cells contacting other N-cadherin^+^ cells and P-cadherin^+^ cells contacting other P-cadherin^+^ cells. The average connection strength between these cells can be quantified by the formula below.

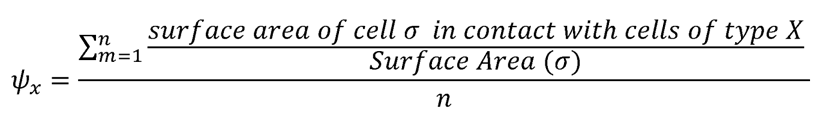

For a cadherin type x (i.e. high E-cadherin), this index calculates how well cells of cadherin type x contact one another. At a given timestep, for a focal cell σ of cadherin type x, if it is in contact with cadherin type x neighbors, its total surface area contact with these cadherin type x neighbors is normalized to the surface area of focal cell σ. This measures the relative strength of cadherin type x connection the cell has to cadherin type x cells at that timestep. Repeating this over all cadherin type x cells in contact with type x cadherin cells and normalizing to the number of contributions yields a global average connection strength for the structure. In this study, as cadherin cells sort to one another, the connection strength increases globally, yielding a measure of sorting. However, this measure can be well generalized depending on the need of the experimenter. See ^33^ for the full theoretical description and capabilities of this index.

### Cell-cell signaling assay quantifications

A frozen 5×5×5 pixel cubic gray cell (i.e. static in volume, surface area, and morphology) was seeded with either 0, 1, 3, or 6 orange 5×5×5 pixel cubic cells such that one face of an orange cell contacted one face of the focal gray cell. This fixed SJR signaling, S_SJR_, to a constant value (i.e. 0 orange cells had L*0/6 S_SJR_ while 1 orange cell had L*1/6 S_SJR_ with L=10000 in this study) and allowed analytically validating the activation circuits as well. For the expression difficulties β=1000, 3000 and 12000, simulations were run for 100000 timesteps with 0, 1, 3, or 6 orange cells deleted at 25000 timesteps. ATF levels were tracked along with red reporter levels. These results were validated by numerically solving the activation circuits in Mathematica via NDSolve, and this enabled generating plots for the additional two expression difficulties shown (β=6000 and 18000). Full parameters are given in Supplementary Table 1.

### 3-layered structure quantifications

A mixture of orange and gray cells carrying the appropriate activation circuit were seeded as a spherical blob at the center of the simulation lattice and simulation run for 50,000 timesteps. The deactivation test (SFig.3B) was run to 75,000 timesteps. Cross section of a 3D structure is shown and scalebar is from the original *in silico* study ^33^, 17.5 pixels to 100 um *in vitro*. Core quality was quantified using the homogeneity index with high E-cadherin^+^ cells (blue/cyan) to other high E-cadherin^+^ cells.

Relative activation was quantified by dividing the number of cells of the focal type over the total cells of that genotype. The formulas are given here. Cyan curve is E-cadherin^+^ cyan cells/ (gray cells + green cells + blue cells + cyan cells). Blue curve is E-cadherin^+^ blue cells/ (gray cells + green cells + blue cells + cyan cells). Green curve is E-cadherin^-^ green cells/ (gray cells + green cells + blue cells + cyan cells). Gray curve is E-cadherin^-^ gray cells/ (gray cells + green cells + blue cells + cyan cells). Full parameters are given in Supplementary Table 1.

### Multipole structure quantifications

A mixture of orange and gray cells carrying the appropriate activation circuit were seeded as a spherical blob at the center of the simulation lattice and simulation run for 50,000 timesteps. Cross section of a 3D structure is shown and scalebar is from the original *in silico* study^33^, 17.5 pixels to 100 um. Pole quality was quantified for N-cadherin^+^ cells to N-cadherin^+^ cells (cyan curves of pole quality in SFig.6) and P-cadherin^+^ cells to P-cadherin^+^ cells (pink curves of pole quality in SFig.6). Relative activation was quantified by the following formulas. Cyan curve is N-cadherin^+^ cyan cells/ (gray cells + green cells + blue cells + cyan cells). Blue curve is N-cadherin^+^ blue cells/ (gray cells + green cells + blue cells + cyan cells). Green curve is N-cadherin^-^ green cells/ (gray cells + green cells + blue cells + cyan cells). Gray curve is N-cadherin^-^ gray cells/ (gray cells + green cells + blue cells + cyan cells). Pink curve is P-cadherin^+^ pink cells/ (orange cells + yellow cells + red cells +pink cells). Red curve is P-cadherin^+^ red cells/ (orange cells + yellow cells + red cells +pink cells). Yellow curve is P-cadherin^-^ yellow cells/ (orange cells + yellow cells + red cells +pink cells). Orange curve is P-cadherin^-^ orange cells/ (orange cells + yellow cells + red cells +pink cells). Full parameters are given in Supplementary Table 1.

### Targeting heterogenous and homogenous tumor quantifications

The appropriate mixture of SB CAR T cells, priming (P) cells, surrounding (S) cells, dual antigen (DA) tumor cells and/or single antigen (U) tumor cells were seeded as a spherical blob at the center of the simulation lattice and simulation run for 50,000 timesteps. Cells were set adhesive to one another (J=35 for all) to prevent random dispersion. CAR levels were tracked and CAR^+^ percentage calculated by (CAR^+^ cyan cells + CAR^+^ blue cells)/ (gray cells + green cells + blue cells + cyan cells). Because orange surrounding (S) cells have the same characteristics as tumor cells, as they are both derived from ISL929, barring antigen expression, they can represent not just normal healthy cells, but also unhindered (not killed by SB CAR T cells) tumor cell growth. Then, tumor cell survival can be calculated at the endpoint with the formula: (dual antigen pink tumor cells + single antigen red tumor cells)/ (n*orange cells), where n=1 when there is only one type of tumor cell and n=2 if both tumor cell types are present. This is effectively normalizing the number of surviving tumor cells to the theoretical endpoint number of tumor cells. This was validated by the control tumor setup: surrounding and tumor cells have similar numbers at the endpoint (SFig.9B). Parameters modified for the challenges are given in the results section and full parameters are given in Supplementary Table 1.

### Statistical analysis

Statistical testing was performed with JMP v11.0.0 with significance level of 0.05. Testing method was decided as planned comparisons before experiments were performed and Wilcoxon test was chosen due to non-parametricity of data. Samples sizes are given in the text or figures or figure captions. I show mean±SEM.

## SUPPLEMENTARY FIGURES

**Supplementary Figure 1.**
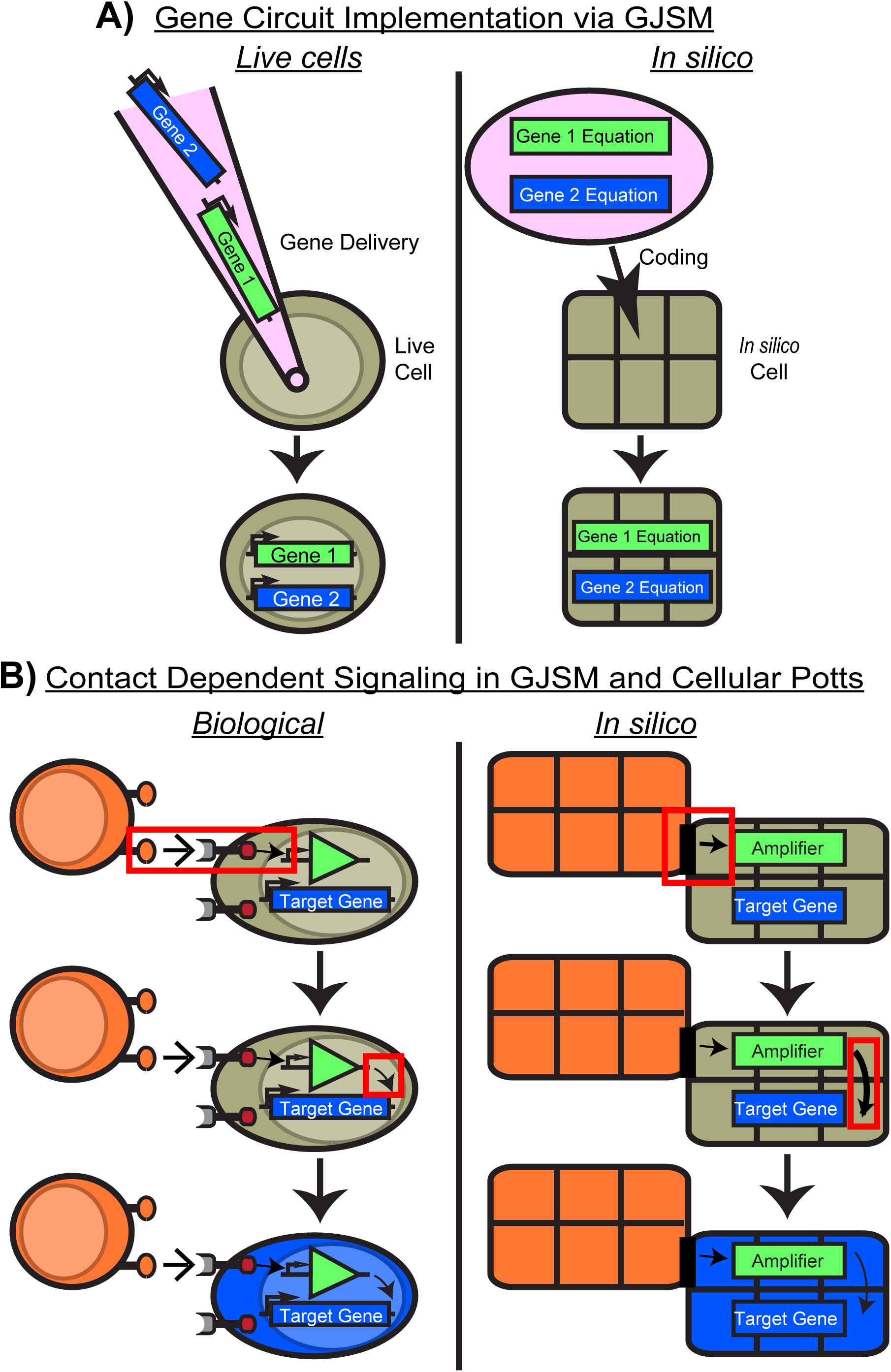
The *In Silico* System is Designed to Parallel the *In Vitro* System, related to. Figure 1. A) The *in silico* (GJSM+CC3D) framework follows the same design principles in engineering biological systems. In place of genetic circuit delivery into live cells, genetic circuits are converted to equations and delivered via coding into i*n silico* cells ^33^. B) An example of the contact dependent SJR signaling process *in silico* compared to biological. I*n silico* juxtacrine signaling mimics biological cell-cell contact dependent SJR signaling by having shared contact pixels between cells dictate the strength of the SJR signal. As in the biological system, this contact dependent signal is then fed through the amplifier and then to the target gene ^33^.

**Supplementary Figure 2.**
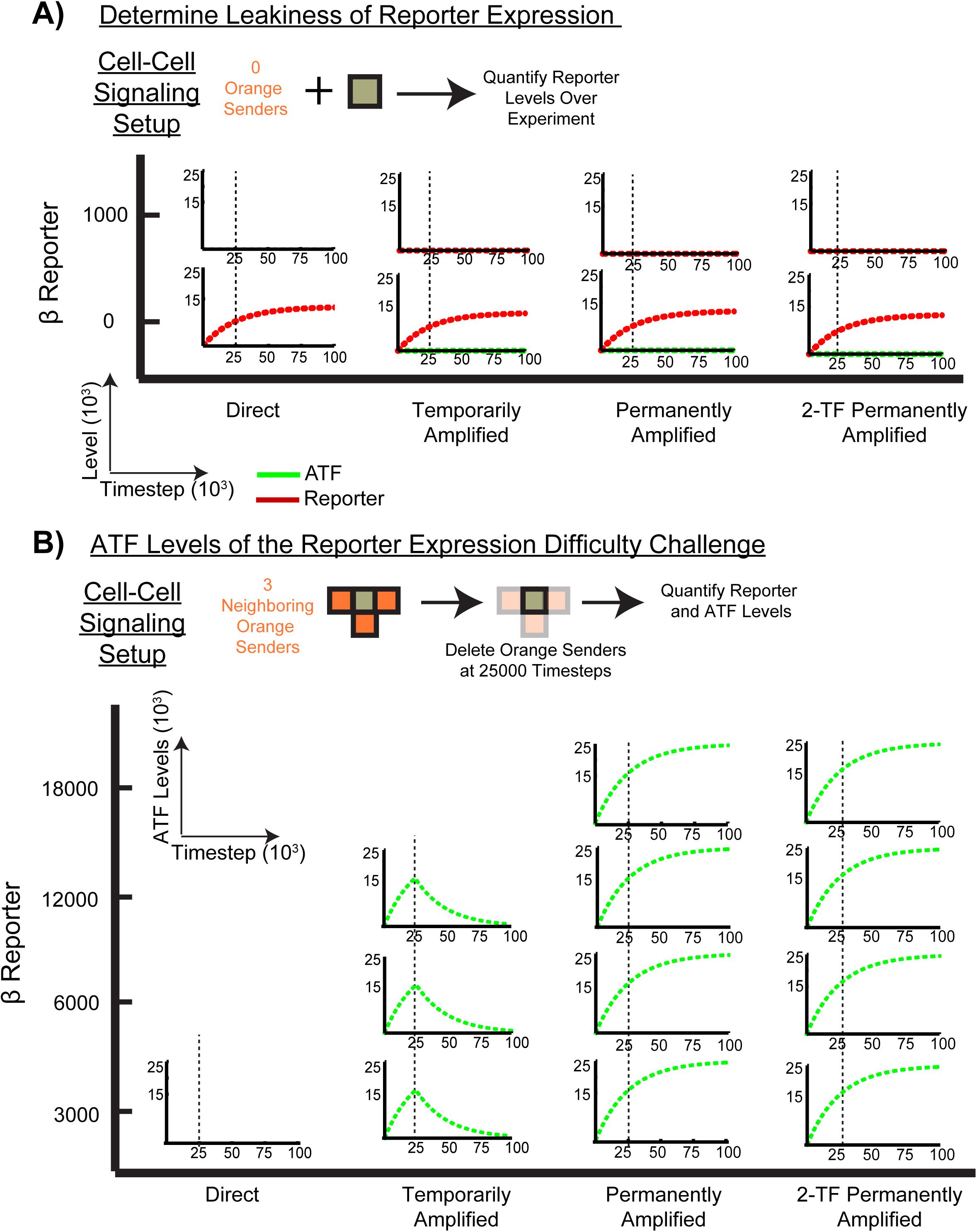
Amplifiers Operate as Designed with the Parameters Tested, related to. Figure 2. A) Uses the same setup as Fig.2A. Gray cells were seeded without orange neighbors to determine leakiness of red reporter expression. At low β Reporter values (i.e. 0), red reporter expression is leaky but at higher values, where the cell-cell signaling simulations are performed, leakiness is not present. Red curve is reporter levels and green line is activating transcription factor (ATF) levels. B) Activating transcription factor (ATF) levels of the different circuits in the reporter expression difficulty challenge. Temporarily amplified activated cells had ATF expression decrease immediately after loss of orange cells while the permanently amplified cells maintained increasing and elevated ATF levels, indicating ATF expression amplification.

**Supplementary Figure 3.**
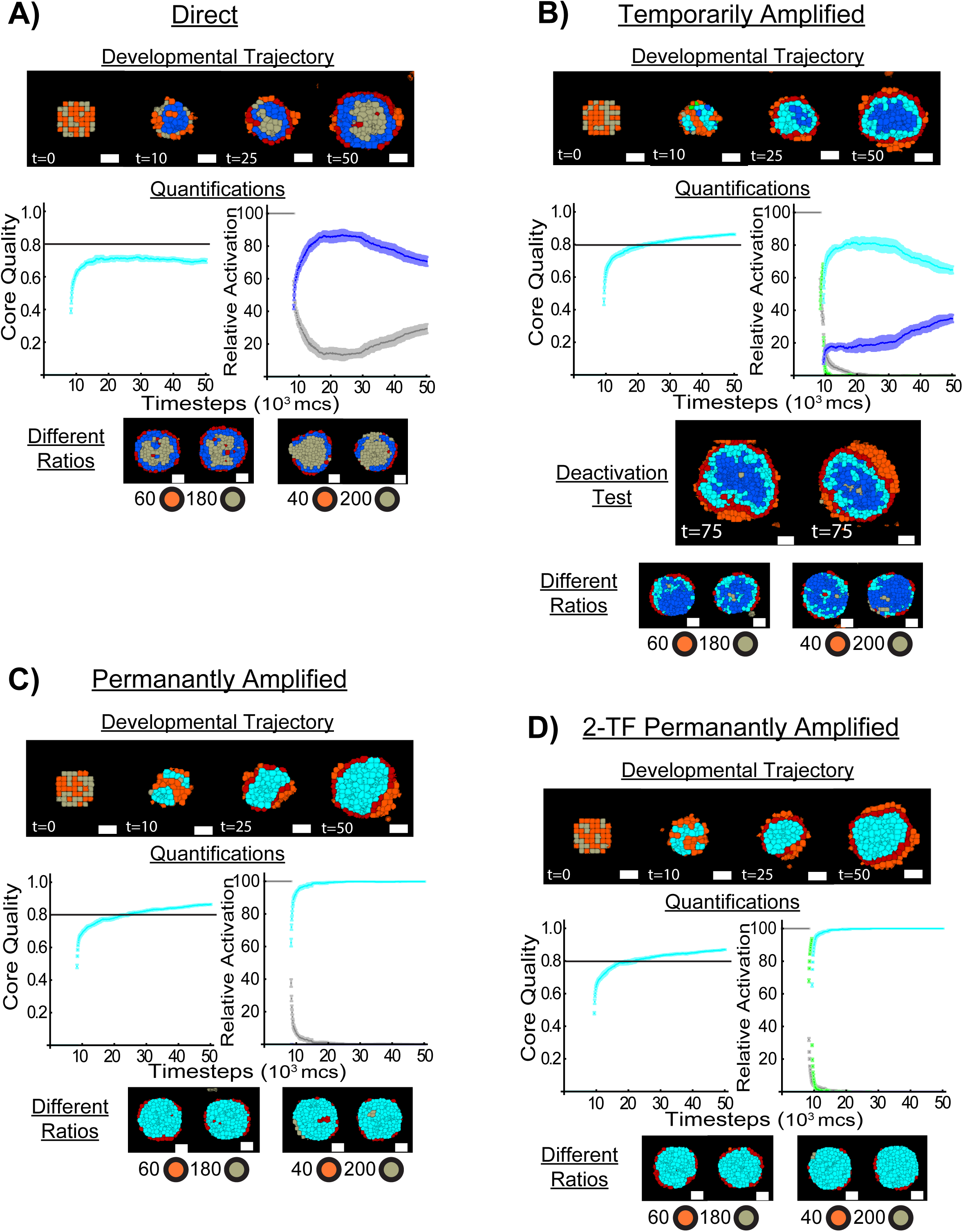
Amplifiers Improve 3-Layered Structure Formation by Amplifying and Prolonging Duration of High E-cadherin and Blue Ligand Expression, related to Figure 3. A) Qualitative and quantitative description of 3-layered structure formation from using directly activated gray cells. In the developmental trajectory (Developmental Trajectory), the majority of gray cells become blue high E-cadherin^+^ but revert as they lose contact with orange cells, resulting in poor core formation. This is reflected quantitatively. Core quality is low <0.8 ^33^ (Core Quality) and the percentage of gray cells that become blue increases but then decreases as time passes (Relative Activation). The gray curve is the percentage of gray cells and the blue curve the percentage of blue cells. Four additional structures (Different Ratios), two per different ratios of orange and gray cells (65±1.22 orange and 186±1.22 gray, 42.60±2.20 orange and 208.40±2.20 gray), at 50,000 timesteps. B) Qualitative and quantitative description of 3-layered structure formation from using temporarily amplified activated gray cells. This amplifier amplifies high E-cadherin/blue ligand expression to allow almost all gray cells become blue or cyan high E-cadherin^+^, resulting in the target 3-layered structure (Developmental Trajectory). This is confirmed quantitatively with the Core Quality and Relative Activation measurements; core quality is higher and more cells are E-cadherin^+^ (blue curve combined with cyan curve) compared to the directly activated cells. Blue, cyan, green, and gray curves correspond to blue, cyan, green, and gray cell percentage, respectively. As the structure develops, however, loss of orange signaling neighbors results in cyan cells (cyan curve) losing ATF expression and becoming blue (blue curve). Nevertheless, they maintain being high E-cadherin^+^ for the duration tested (50,000 timesteps). Extending the simulation time to 75,000 timesteps shows that blue cells begin reverting to gray, confirming that the temporarily amplified activation circuit only temporarily prolongs high E-cadherin and blue ligand expression (Deactivation Test). Four additional structures (Different Ratios), two per different ratios of orange and gray cells (65±1.22 orange and 186±1.22 gray, 42.60±2.20 orange and 208.40±2.20 gray), at 50,000 timesteps. C) Qualitative and quantitative description of 3-layered structure formation from using permanently amplified gray cells. Developmental trajectory shows that the amplifier amplifies and prolongs high E-cadherin/blue ligand expression such that nearly all gray cells become high E-cadherin^+^ (blue or cyan) to robustly form the 3-layered structure. This is confirmed quantitatively with the higher core quality (Core Quality) and higher percentage of high E-cadherin^+^ cells that do not revert (blue and cyan curves in Relative Activation) compared to the directly activated cells. Blue, cyan, green, and gray curves correspond to blue, cyan, green, and gray cell percentage, respectively. Four additional structures (Different Ratios), two per different ratios of orange and gray cells (65±1.22 orange and 186±1.22 gray, 42.60±2.20 orange and 208.40±2.20 gray), at 50,000 timesteps. D) Qualitative and quantitative description of 3-layered structure formation from using 2-TF permanently amplified activated gray cells. Results are similar qualitatively and quantitatively to SFig.3C. The amplifier amplifies high E-cadherin/blue ligand expression to convert almost all gray cells to cyan cells and demonstrates no reversion of blue/cyan cells, confirming its permanency of high E-cadherin/blue ligand expression. Four additional structures (Different Ratios), two per different ratios of orange and gray cells (65±1.22 orange and 186±1.22 gray, 42.60±2.20 orange and 208.40±2.20 gray), at 50,000 timesteps. N=5 for each set of parameters. Simulations run for 50,000 timesteps.

**Supplementary Figure 4.**
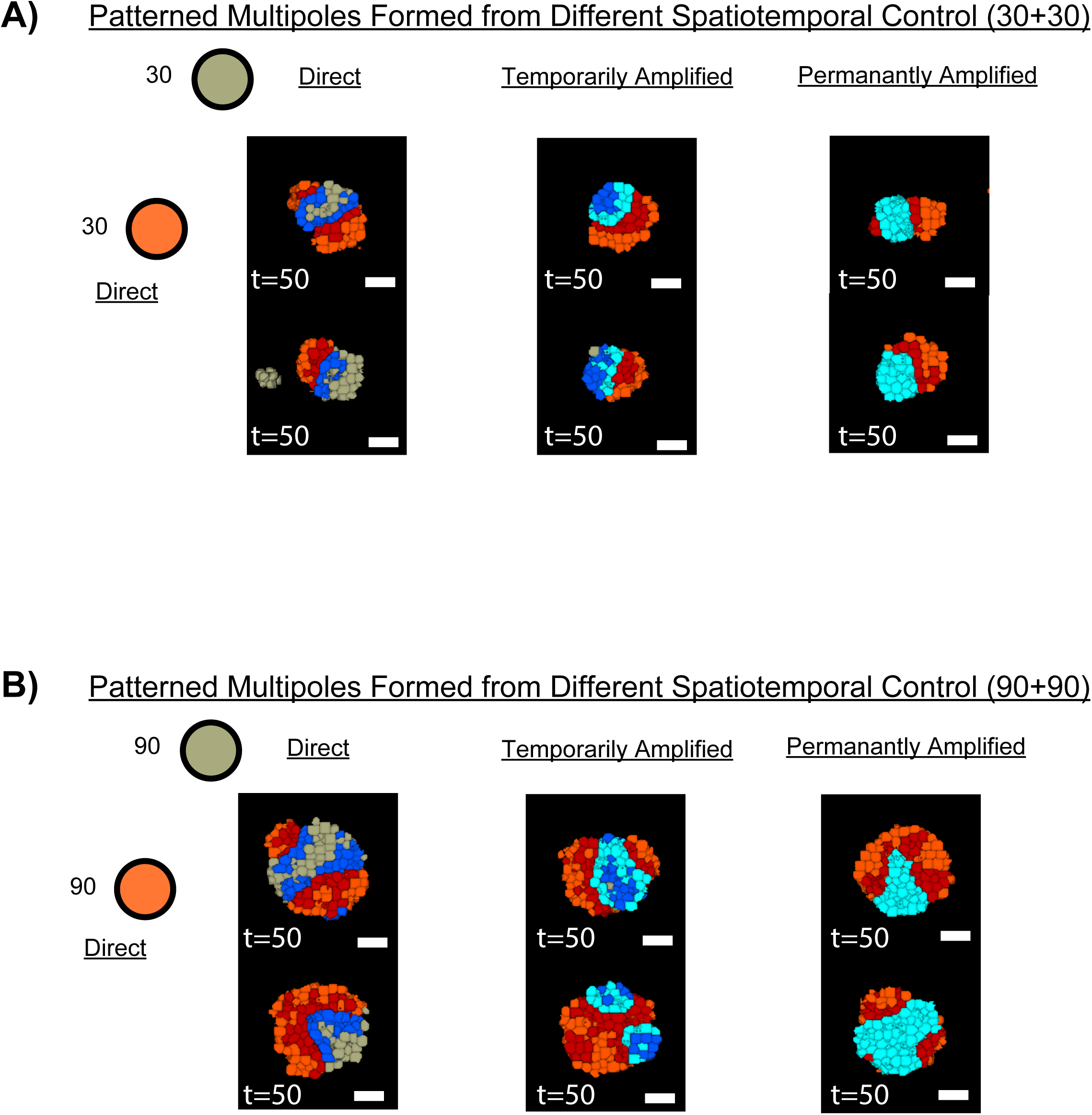
Additional Multipoles Structures Formed from Different Spatiotemporal Control from Different SJR Amplifier Circuits, related to Figure 4. A) Additional structures from combining directly activated orange cells with differently activated gray cells. Different amplifiers enable different temporal control that dictates spatiotemporal patterning. Mixtures are of 27.4±1.04 orange cells with 29.6±1.04 gray cells. N=10 for each circuit combination with two representative cross sections shown. Simulations run for 50,000 timesteps. B) Additional structures from combining directly activated orange cells with differently activated gray cells. Mixtures are of 87.4±1.47 orange cells with 91.6±1.47 gray cells. N=10 for each circuit combination with two representative cross sections shown. Simulations run for 50,000 timesteps.

**Supplementary Figure 5.**
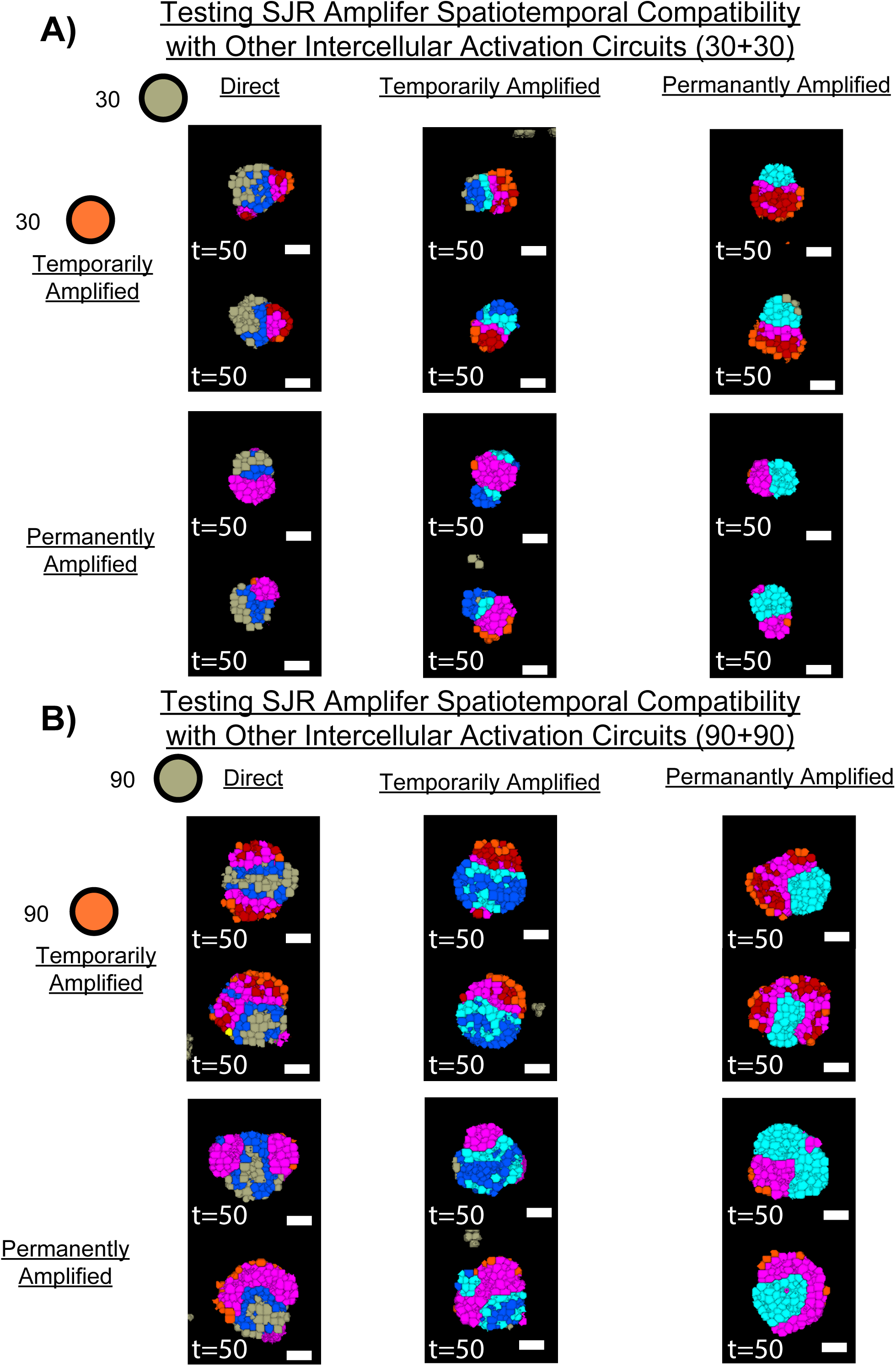
Additional Structures from Testing the Intercellular Compatibility of the Spatiotemporal Control from SJR Amplifier Circuits, related to Figure 5. A) Additional structures from combining differently activated orange cells with differently activated gray cells. Orange cells spatiotemporally pattern independent of the activation circuit in gray cells. Mixtures are of 27.4±1.04 orange cells with 29.6±1.04 gray cells. N=10 for each circuit combination with two representative cross sections shown. Simulations run for 50,000 timesteps. B) Additional structures from combining different activated orange cells with differently activated gray cells. Mixtures are of 87.4±1.47 orange cells with 91.6±1.47 gray cells. N=10 for each circuit combination with two representative cross sections shown. Simulations run for 50,000 timesteps.

**Supplementary Figure 6.**
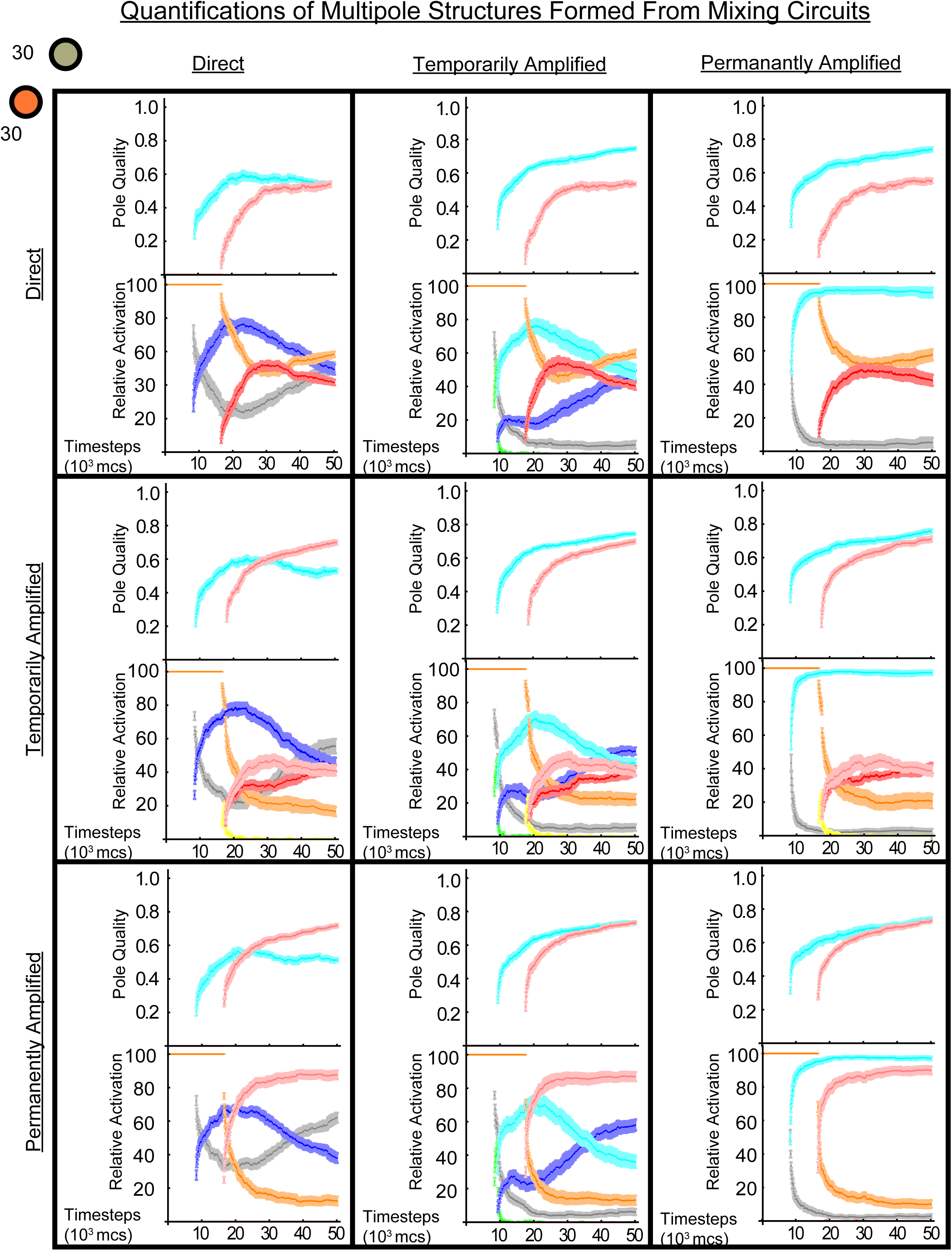
Quantifications of Multipole Structures Formed by Mixing Differently Activated Orange Cells with Differently Activated Gray Cells, related to Figure 4 **and 5**. Pole quality of all the shown multipole structures from testing both spatiotemporal control (Fig.4) and spatiotemporal intercellular compatibility (Fig.5) is given with cyan curve calculated from blue/cyan N-cadherin^+^ cells contacting N-cadherin^+^ cells and pink curve calculated from red/pink P-cadherin^+^ cells contacting P-cadherin^+^ cells. See Homogeneity Index in the Methods Section for more information. Relative activation of all the resulting multipole structures is given as well. Curve color corresponds to cell state color from Fig.5B (i.e. cyan curve represents cyan cells) and is normalized to cell genotype (gray, green, blue, and cyan are one genotype and orange, yellow, red, and pink are one genotype). See Multipole Structure Quantifications in the Methods Section for more details. Mixtures are of 27.4±1.04 orange cells with 29.6±1.04 gray cells. Curves from n=10 for each circuit combination.

**Supplementary Figure 7.**
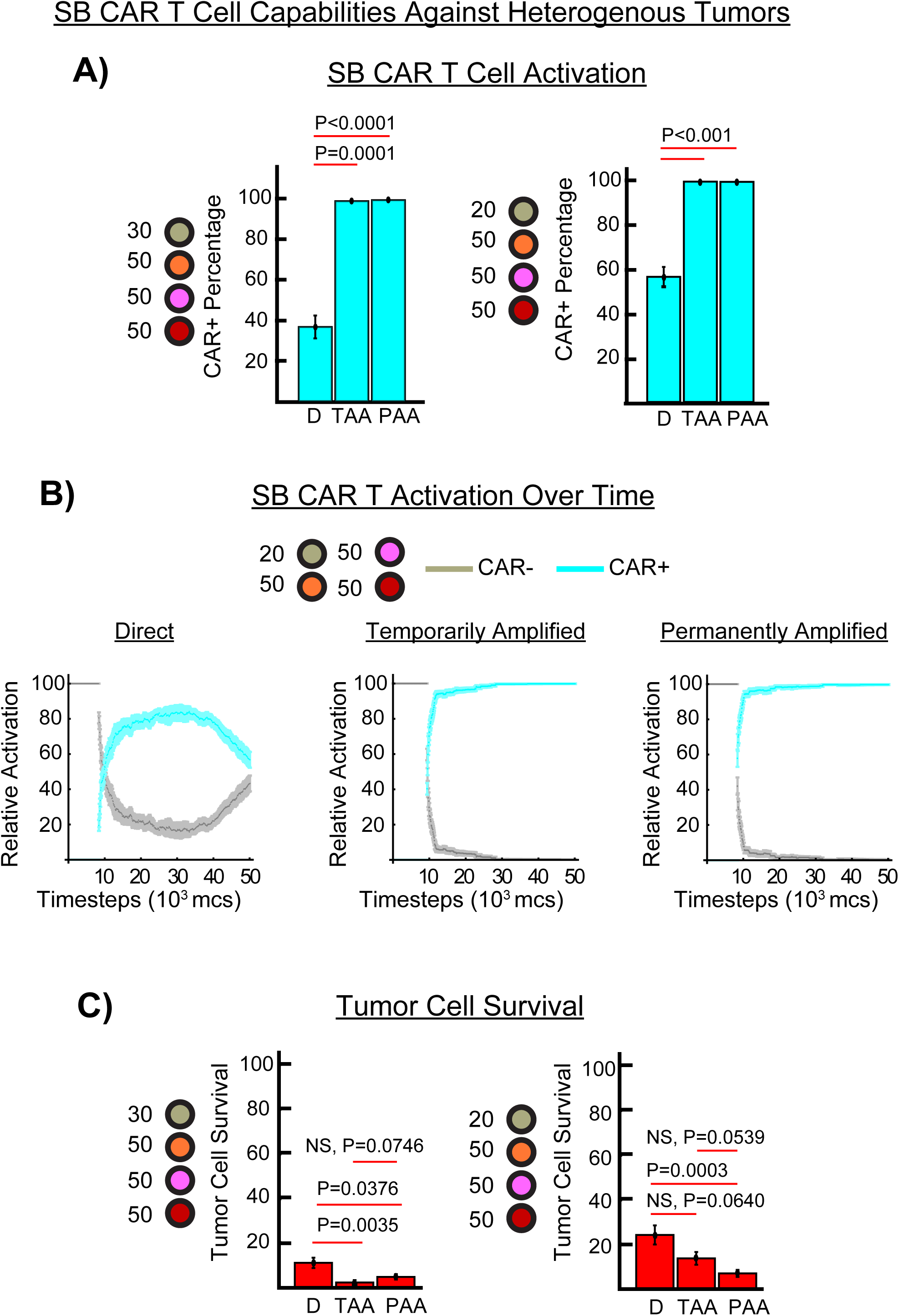
Amplifier Imbued SB CAR T Cell Capabilities Against Heterogeneous Tumors Under Optimal Conditions, related to Figure 6. A) Percentage of CAR^+^ cells against two different heterogenous tumors (27.8±1.44 gray SB CAR T cells with 47.1±1.13 orange S cells with 49.6±1.70 red U cells and 54.5±1.42 pink DA cells) and (19.8±0.85 gray SB CAR T cells with 55.7±2.13 orange S cells with 52.5±2.33 red U cells and 51±1.82 pink DA cells) at the 50,000 timesteps endpoint. D is direct activation, TAA is temporarily amplified activation, and PAA is permanently amplified activation. B) SB CAR T cell activation to CAR^+^ over time in the latter heterogenous tumor setup. Amplifiers enabled more cells to become CAR^+^ (cyan curve) and maintain being CAR^+^, confirming the amplification and temporal features of the amplifiers. Gray curve is CAR^-^ percentage. C) Tumor cell killing by the SB CAR T cells. Amplifiers overall enabled cells to kill more tumor cells. N=10 for each circuit in each heterogenous tumor. Significance and p value is given. Simulations run for 50,000 timesteps.

**Supplementary Figure 8.**
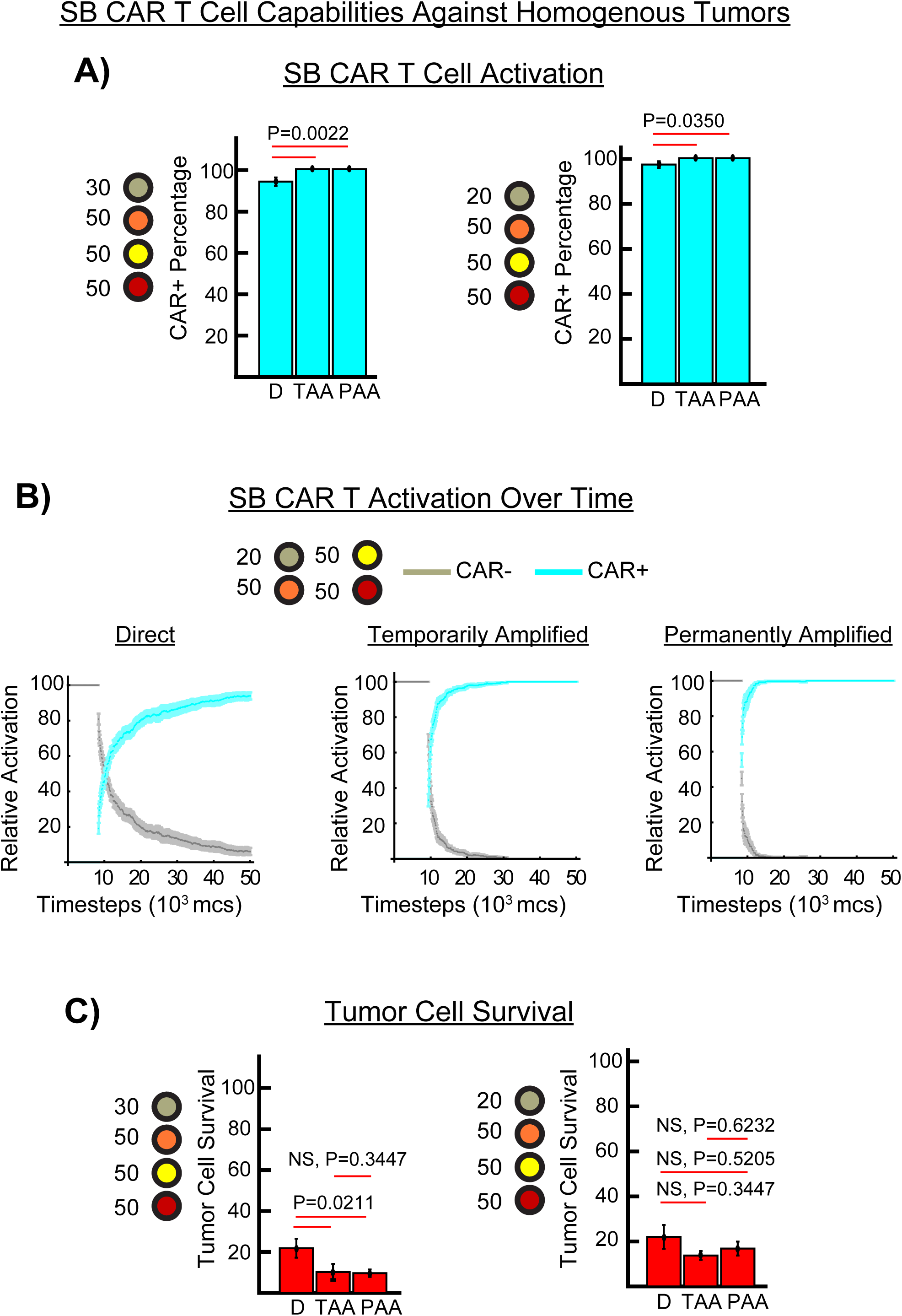
Amplifier Imbued SB CAR T Cell Capabilities Against Homogenous Tumors Under Optimal Conditions, related to. Figure 7. A) Percentage of CAR^+^ cells against two different homogenous tumors (27.8±1.44 gray SB CAR T cells with 47.1±1.13 orange S cells with 49.6±1.70 yellow P cells and 54.5±1.42 red U cells) and (19.8±0.85 gray SB CAR T cells with 55.7±2.13 orange S cells with 52.5±2.33 yellow P cells and 51±1.82 red U cells) at the 50,000 timesteps endpoint. D is direct activation, TAA is temporarily amplified activation, and PAA is permanently amplified activation. B) SB CAR T cell activation to CAR^+^ over time in the latter homogenous tumor setup. Amplifiers in general enabled more cells to become CAR^+^ (cyan curve) earlier than did the direct activation circuit but at the endpoint all circuits have most cells being CAR^+^. Gray curve is CAR^-^ percentage. C) Tumor cell killing by the SB CAR T cells. Amplifier imbued SB CAR T cells appeared to kill more tumor cells and was significant in homogenous tumor with ∼30 SB CAR T cells and ∼50 of the other cells. N=10 for each circuit in each homogeneous tumor. Significant difference and p value is given. Simulations run for 50,000 timesteps.

**Supplementary Figure 9.**
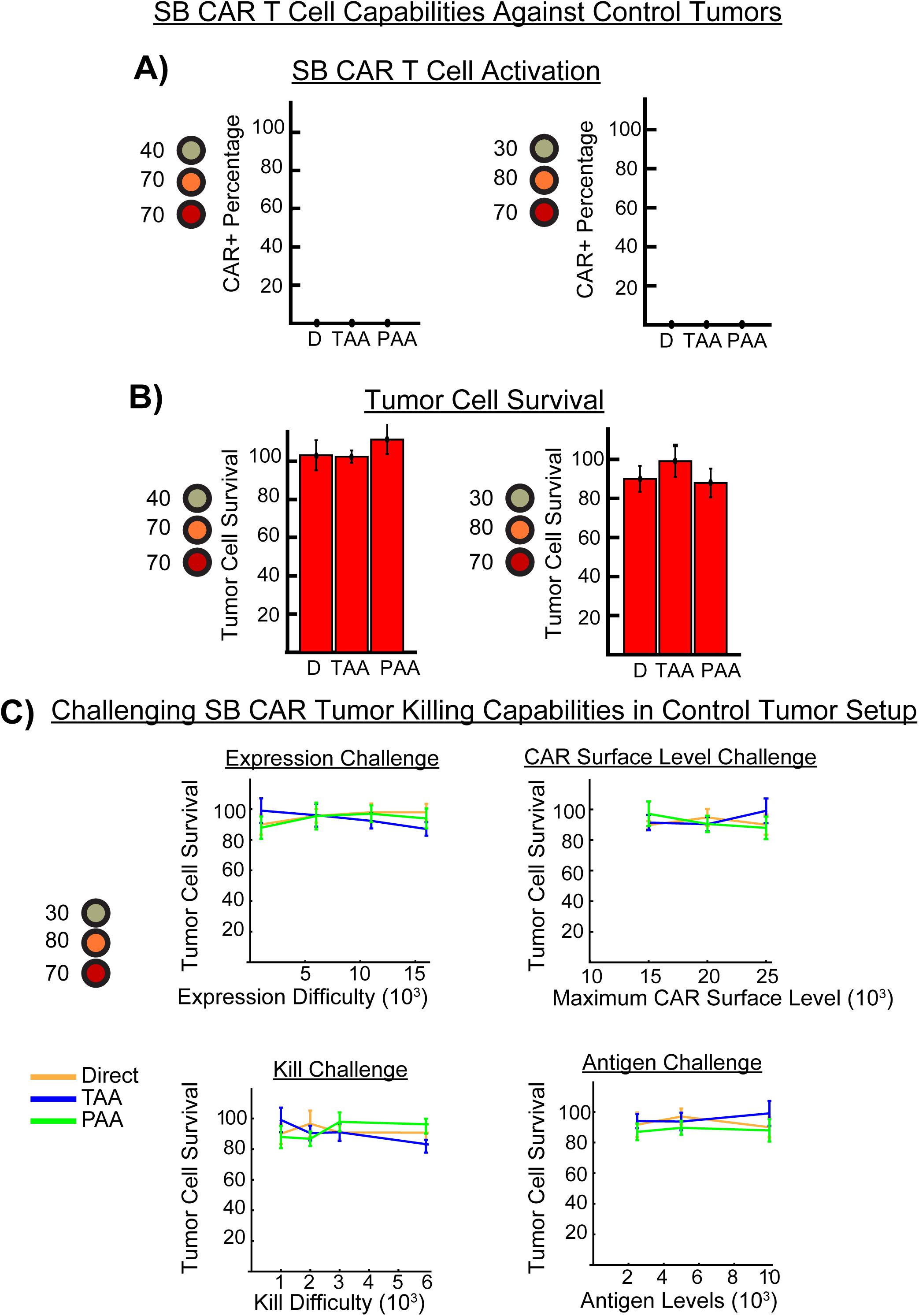
Amplifier Imbued SB CAR T Cells Require Tumor Antigen One/Priming Antigen to Kill Tumor Cells, related to. Figures 6 and 7. A) Percentage of CAR^+^ cells against two control tumors with no tumor antigen one or priming antigen (36.9±1.74 gray SB CAR T cells with 69.7±1.86 orange S cells and 72.4±1.69 red U cells or 27.8±1.44 gray SB CAR T cells with 79±1.47 orange S cells and 72.2.5±1.83 red U cells). Lack of required antigen prevented SB CAR T cells from becoming CAR^+^. D is direct activation, TAA is temporarily amplified activation, and PAA is permanently amplified activation. B) Percentage of tumor cell survival in both setups. Lack of required antigen prevented SB CAR T cells from killing tumor cells. C) SB CAR T cells were subjected to the same challenges as in the heterogenous and homogenous tumor, but lack of the required antigen prevented any tumor cell killing. No significant difference was detected. N=10 for each circuit for each parameter tested. * denotes significant difference and is colored the curve it differs from. Simulations run for 50,000 timesteps.

